# Red light controls adventitious root regeneration by modulating hormone homeostasis in *Picea abies* seedlings

**DOI:** 10.1101/2020.03.11.985838

**Authors:** Sanaria Alallaq, Alok Ranjan, Federica Brunoni, Ondřej Novák, Abdellah Lakehal, Catherine Bellini

## Abstract

Vegetative propagation relies on the capacity of plants to regenerate *de novo* adventitious roots (ARs), a quantitative trait controlled by the interaction of endogenous factors such as hormones and environmental cues among which light plays a central role. However, the physiological and molecular components mediating light cues during AR initiation (ARI) remain largely elusive. We explored the effect of light spectral quality on ARI in de-rooted Norway spruce seedlings as well as on hormone metabolism with sensitive mass spectrometry-based methods. We coupled this to gene expression analysis to identify potential signaling pathways and to extensive anatomical characterization to investigate ARI at the cellular level. We showed that in contrast to white light and blue light, red light promoted ARI likely by reducing jasmonate (JA) and JA-isoleucine biosynthesis and repressing the accumulation of isopentyl-adenine-type cytokinins and abscisic acid. We confirmed that exogenously applied JA and/or CK inhibit ARI, and found that they possibly act in the same pathway. The negative effect of JA was confirmed at the histological level. We showed that JA represses the early events of ARI. In conclusion, RL promotes ARI by repressing the accumulation of the wound-induced phytohormones JA and CK.

**Highlight:** Blue and red light have an opposite effect on adventitious root initiation in Norway spruce hypocotyl, red light having a promoting effect by modulating hormone homeostasis.

## Introduction

Vegetative (or clonal) propagation through stem cuttings is widely used in breeding programs for rapid fixation of elite genotypes. Successful clonal propagation requires *de novo* regeneration of adventitious roots (ARs), a process governed by a multitude of developmentally-programmed and environmentally-induced cues including wounding, light and nutrient availability (Bellini *et al*., 2014; Steffens & Rasmussen, 2016). Because AR regeneration is a quantitative trait, the rooting capacity of cuttings varies among individuals within species, populations, or even clones (Abarca & Díaz-Sala, 2009). Phytohormones, mainly auxin, jasmonate (JA), cytokinins (CKs), ethylene and abscisic acid (ABA), play major roles in AR formation. It has been shown that there is interplay among these phytohormones in complex networks and loops that ensure proper AR development (Lakehal & Bellini, 2019). For example, in *Arabidopsis thaliana*, it has been shown that auxin signaling controls AR formation by modulating JA homeostasis in the hypocotyl (Gutierrez *et al*., 2012; Lakehal *et al*., 2019a) and that JA cooperates with CKs to repress AR formation (Lakehal *et al*., 2020a). Moreover, JA signaling induces the expression of *DIOXYGENASE FOR AUXIN OXIDATION 1*, a key enzyme catalysing the irreversible degradation of auxin, to ensure robust crosstalk between auxin and JA signaling pathways (Lakehal *et al*., 2019b). However, the extent to which these complex molecular networks are evolutionarily conserved among plant species is poorly understood, especially for trees. Although it remains difficult to address the molecular mechanisms controlling AR formation in perennial species, the rapid development of sequencing technologies giving access to most genome and transcriptome sequences now offers the possibility of further exploring the degree of evolutionary conservation or divergence of the known molecular networks.

Light is one of the most important external cues controlling plant growth and development. It is perceived by highly sophisticated sensing machinery which mediates a plethora of physiological and developmental processes, *inter alia* photosynthesis, photomorphogenesis, flowering time and tropisms (de Wit *et al*., 2016). Depending on light intensity and spectral quality, the cues perceived trigger specific downstream transcriptional signatures enabling a plant to adjust its growth and development as well as its response to the environment. Phytohormones play central roles in integrating and translating cues into developmentally coherent outputs and responses (de Wit *et al.*, 2016). In particular, light intensity and spectral quality have long been manipulated as tools in vegetative propagation practices in order to stimulate the rooting ability of recalcitrant species. For example, treating donor plants of difficult-to-root *Eucalyptus globulus* species with far-red-enriched light stimulated the rooting capacity of microcuttings (Ruedell *et al*., 2015). This stimulation was possibly due to upregulation of the expression of *AUXIN RESPONSE FACTOR6-like (ARF6-like)* and *ARF8-like* genes at the cutting site by far-red light. These transcriptional regulators have been shown to promote ARI in Arabidopsis hypocotyls (Gutierrez *et al*., 2009, 2012) and in *Populus* stem cuttings (Cai *et al*., 2019). Far-red-enriched light also induced the expression of a putative indole 3-acetic acid (IAA) biosynthetic gene, *YUCCA3-like*, and two putative auxin efflux carriers genes, *PIN-FORMED 1-like (PIN1-like)* and *PIN-FORMED 2-like (PIN2-like)*, in *Eucalyptus globulus* (Ruedell *et al*., 2015). These data suggest that far-red-enriched light might act on the auxin biosynthesis, transport and/or signaling pathways to stimulate AR formation in *Eucalyptus globulus.* Keeping leafy stem cuttings of *Petunia hybrida* (Klopotek *et al.*, 2016) or *Dianthus caryophyllus* (Agulló-Antón *et al*., 2011) in dark conditions prior to light exposure improved their rooting capacity. The effect of light on Norway spruce (*Picea abies*) growth and morphogenesis has also been documented (Strömquist & Eliasson, 1979; Mortensen & Sandvik, 1988; Bollmark & Eliasson, 1990; Mølmann *et al*., 2006; Burescu *et al*., 2015; OuYang *et al*., 2015) and it has been shown that high-irradiance white light inhibited AR in Norway spruce stem cuttings (Strömquist & Eliasson, 1979).

Despite the importance of light in regulating AR formation, the physiological and molecular mechanisms involved are still unknown. Here, we used de-rooted Norway spruce seedlings as a model system to explore the role of light in the control of AR initiation in conifers. We found that constant red light promotes this process, possibly through downregulation of JA signaling and repression of cytokinin and abscisic acid (ABA) accumulation compared to constant white light conditions. It appears that the cooperative role of JA and CK signaling in the repression of ARI is evolutionarily conserved, as it represses very early events in the ARI process in Norway spruce hypocotyls.

## Materials and methods

### Plant material

Commercial seeds from Norway spruce (*Picea abies* L. Karst) were provided by Svenska Skogsplantor (https://www.skogsplantor.se/; Sweden) and kept at a temperature of 4 °C prior to use. The seeds were soaked in tap water for 24 h, in the dark at 4 °C. They were then sown in fine wet vermiculite as described in Ricci *et al.* (2008) and allowed to germinate and grow in a growth chamber. The seedlings were watered twice a week with tap water and left to grow for three weeks (Supplementary Fig. S1A). The growth conditions were as follow: 16 h light with 75 μmol /m^2^/s (400 to 700 nm, Fig. S2A); 8 h dark; 20°C day temperature and 18°C night temperature. The white light was provided by 36W/840 Super 80 Cool White TL-D tubes (Philips, https://www.philips.se/).

### Test of effects of different light conditions on adventitious rooting of de-rooted seedlings

Twenty-one days after sowing, the seedling hypocotyls were severed 2 cm below the apex (Supplementary Fig. S1B) and placed in 24 ml vials filled with distilled water (Supplementary Fig. S1C). Three vials containing 5 seedlings each, constituting technical replicates, were prepared.

In order to test different white light conditions, vials containing de-rooted seedlings were kept in either long-day conditions in the growth chamber as described above (Supplementary Fig. S2A) or in a Percival growth cabinet equipped with cool white fluorescent tubes (F17T8/TL741, 17 Watt; Philips U.S.A: Percival Scientific, Perry Iowa, USA) (Supplementary Fig. S2B) or in a Percival plant growth cabinet (E-*30NL/floraLED*) equipped with six CLF *floraLED* modules (CLF PlantClimatics GmbH; Emersacker, Germany) providing white, red or blue light (Supplementary Fig. S2C,E,F). Under these latter conditions the light was constant and referred to as <u>c</u>onstant <u>W</u>hite <u>L</u>ight (cWL), <u>c</u>onstant <u>B</u>lue <u>L</u>ight (cBL) or <u>c</u>onstant <u>R</u>ed <u>L</u>ight (cRL), and the temperature was constant and set at 20 °C.

In order to remove the blue region of the spectrum from the white LEDs we added a yellow filter, which almost totally abolished the blue peak at 460 nm (Supplementary Fig. S2D).

The cBL (460 nm) intensity was set at 9 μmol/m^2^/s (Supplementary Fig. S2E), and cRL (660 nm) was set at an intensity of 9 μmol /m^2^/s (Supplementary Fig. S2F).

The vials were refilled daily with distilled water to replace the water lost by evapotranspiration. The number of roots per cutting and the number of rooted cuttings were monitored for 30 days after cutting (DAC) and all the experiments were repeated at least three times.

### Hormone treatments

Auxin treatment:

Five hypocotyl cuttings per vial, with three technical replicates, were cultured in 24 ml of distilled water supplemented with indole butyric acid (IBA), indole acetic acid (IAA) or 1-Naphthalene Acetic Acid (NAA) at final concentrations of 1 or 5 μM, or without supplementation as mock treatment.

Jasmonate (JA) treatment:

Five hypocotyl cuttings per vial, with three technical replicates, were cultured in 24 ml of distilled water supplemented with JA at final concentrations of 2 μM, 10 μM or 20 μM, or without supplementation as mock treatment.

Cytokinin treatment:

Five hypocotyl cuttings per vial, with three technical replicates, were cultured in 24 ml of distilled water supplemented with 6-benzylaminopurine (BAP) at a final concentration of 0.1 μM or 0.01 μM, or without supplementation.

BAP was also used in combination with 1 μM IBA, 2 μM or 5 μM JA.

In all these experiments vials were covered with parafilm in order to limit the evaporation of hormone solutions during the rooting period; small holes allowed the cuttings to be inserted into the vials.

### Histology

Five mm sections of hypocotyl cuttings were dissected from three-week-old seedlings at T0, just before transferring the cuttings to rooting conditions, then 3, 5, 10, 13 and 15 days after cutting. Similar material was taken from de-rooted seedlings treated with 1 μM IBA or 20 μM JA. Hypocotyl samples were vacuum infiltrated with fixation medium, 37% formalin, 5% acetic acid, 100% ethanol, for 20 seconds and left for 24 hours at room temperature. The samples were then washed in fixation solvent (70 % ethanol) for 10 minutes and transferred into 70 % EtOH until required for use. Samples were then gradually dehydrated in an ethanol series (80 %, 90 %, 96 % for 2h each and 100 % overnight at room temperature). The 100 % EtOH was gradually replaced by HistoChoice tissue fixative (VWR Life, https://us.vwr.com/) in three steps of 1:3, 1:1, 3:1 (EtOH: HistoChoice ratio) and two 1 h steps with pure HistoChoice. The HistoChoice fixative was gradually replaced with Paraplast Plus for tissue embedding (Sigma-Aldrich, https://www.sigmaaldrich.com/), over 6 days. Serial sections (10 μm) were obtained with a Zeiss HM 350 rotary microtome (https://www.zeiss.com/). Longitudinal and transverse sections were stained with safranin (Sigma-Aldrich, https://www.sigmaaldrich.com/) and alcian blue (Sigma-Aldrich, https://www.sigmaaldrich.com/) in a ratio of 1:2; using methods from (Hamann *et al*., 2011).

### Sources of sequence data and phylogenetic tree construction

The *AtMYC2* (At1g32640), *AtJAZ3* (At3g17860), *AtJAZ10* (At5g13220), *AtAOC1* (At3g25760) and *AtCOI1* (At2g39940) *Arabidopsis thaliana* gene coding sequences (CDS) from TAIR (https://www.arabidopsis.org) were used as queries in BLASTN searches for CDS predicted with high confidence from *P. abies* (http://congenie.org/; Nystedt *et al*., 2013) to identify potential orthologs in the *Picea abies* genome. All sequence information is provided in Supplementary Fig. S4. To find the putative Arabidopsis orthologs in *Picea abies*, a BLAST alignment of the coding sequences of selected genes was generated using “Genome tools” from the Congenie database with default settings. Subsequently, the sequences were used to construct phylogenetic trees with the tools available at “http://www.phylogeny.fr”. The phylogenetic analysis was based on the neighbor-joining method (with 1000 bootstrap replicates). The putative orthologs were named according to the Congenie database.

### Primer design, efficiency and qRT-PCR analysis

Candidate genes were identified by BLASTN searches against the spruce database (http://congenie.org) using Arabidopsis protein coding nucleotide sequences. The Norway spruce genes that showed the highest sequence similarity with Arabidopsis genes were further considered for primer design and gene expression analysis. Primers for the selected candidate genes and reference genes to be used for qRT-PCR were designed using Primer3web version 4.1.0 (http://primer3.ut.ee); the primer sequences are listed in Table **S1**. Specificity of the primers was analyzed by Sanger sequencing of amplified PCR products and cDNA was used as a template for PCR amplification from each primer pair. For each primer pair, PCR reaction efficiency estimates were calculated from a standard curve generated from a serial dilution of cDNA. Based on the average cycle threshold (Ct) values for each five-fold dilution, a standard curve was generated using linear regression. PCR efficiency (E) was derived from the equation: Efficiency % = (10 (−1/slope) - 1) ×100%. PCR conditions were as follows: 95°C for 1 min, 40 cycles of 95°C for 15 s, 60°C for 30 s, and 72°C for 30 s. Finally, dissociation curves were generated by increasing the temperature from 65 to 95°C, stepwise by 0.3°C every 10 s. For gene expression analysis by quantitative PCR (qPCR), total RNA was isolated from the base of spruce hypocotyls (5 mm long from 36 to 38 hypocotyl cuttings) using a Spectrum™ Plant Total RNA Kit (Sigma-Aldrich; https://www.sigmaaldrich.com/). cDNA was synthesized using an iScript™ cDNA Synthesis Kit (Bio-Rad; https://www.bio-rad.com/en-se/) following DNase treatment. Expression was quantified by the LightCycler^®^ 480 System (Roche, Basel, Switzerland) using LightCycler^®^ 480 SYBR Green I Master (Roche, Basel, Switzerland). For each sample, three technical replicates from two independent biological replicates were analyzed. The ΔΔ^CT^ method was used to measure the relative amount of transcript from each gene. Expression values were normalized to the expression of two reference genes, *PaEIF4A1* and *PaUBC28*, and calibrated with the control value arbitrarily set to 1.

In order to check whether expression of the selected putative orthologs *PaMYC2-like, PaAOC1-like, PaJAZ3-like* and *PaJAZ10-like* could be induced by jasmonate, 3-week-old whole Norway spruce seedlings were treated with 50 μM of JA or mock treatment for 3 h. Total RNA was prepared and qPCR experiments performed as described above.

### Hormone profiling experiments

For hormone quantification, 5 mm hypocotyl segments were dissected at the base of de-rooted hypocotyls 6, 24, 48, 72 and 120 hours after they had been transferred to cWL, cBL or cRL, in order to provide an average of 20 mg of fresh weight.

Quantification of cytokinin metabolites was performed according to the method described by (Svačinová et al., 2012), including modifications described by (Antoniadi et al., 2015). Samples (20 mg fresh weight, FW) were homogenized and extracted in 1 ml of modified Bieleski buffer (60% MeOH, 10% HCOOH and 30% H_2_O) together with a cocktail of stable isotope-labeled internal standards (0.25 pmol of CK bases added per sample). The extracts were applied **to** an Oasis MCX column (30 mg/1 ml, Waters) conditioned with 1 ml each of 100% MeOH and H_2_O, equilibrated sequentially with 1 ml of 50% (v/v) nitric acid, 1 ml of H_2_O, and 1 ml of 1M HCOOH, and washed with 1 ml of 1M HCOOH and 1 ml 100% MeOH. Analytes were then eluted in two steps using 1 ml of 0.35M NH4OH aqueous solution and 2 ml of 0.35M NH4OH in 60% (v/v) MeOH. The eluates were then evaporated to dryness *in vacuo* and stored at −20°C. Cytokinin levels were determined by ultra-high performance liquid chromatography-electrospray tandem mass spectrometry (UHPLC-MS/MS) using stable isotope-labeled internal standards as a reference (Rittenberg and Foster, 1940). Separation was performed on an Acquity UPLC^®^ System (Waters, Milford, MA, USA) equipped with an Acquity UPLC BEH Shield RP18 column (150×2.1 mm, 1.7 μm; Waters), and the effluent was introduced into the electrospray ion source of a triple quadrupole mass spectrometer Xevo™ TQ-S MS (Waters). Six independent biological replicates were performed.

Endogenous levels of free IAA, ABA, SA and JA, as well as the conjugated form of JA, JA-Ile, in 20 mg of hypocotyls were determined according to the method described in (Floková *et al*., 2014). The phytohormones were extracted using an aqueous solution of methanol (10% MeOH/H_2_O, v/v). To validate the LC-MS method, a cocktail of stable isotope-labeled standards was added with the following composition: 5 pmol of [^13^C_6_]IAA, 10 pmol of [^2^H_6_]JA, [^2^H_2_]JA-Ile and 20 pmol of [^2^H_4_]SA (all from Olchemim Ltd, Czech Republic) per sample. The extracts were purified using Oasis HLB columns (30 mg/1 ml, Waters, Milford, MA, USA) and targeted analytes were eluted using 80% MeOH. Eluate containing neutral and acidic compounds was gently evaporated to dryness under a stream of nitrogen. Separation was performed on an Acquity UPLC^®^ System (Waters, Milford, MA, USA) equipped with an Acquity UPLC BEH C18 column (100 x 2.1 mm, 1.7 μm; Waters, Milford, MA, USA), and the effluent was introduced into the electrospray ion source of a triple quadrupole mass spectrometer Xevo^™^ TQ-S MS (Waters, Milford, MA, USA).

### Statistical analysis

For statistical analyses, the SPSS 20 software package was used (https://www.ibm.com/support/pages/spss-statistics-20). Student’s t-tests were performed to evaluate the effects of different light qualities on hormone contents.

Repeated measures ANOVAs were used to test the effects of time and hormone concentrations under constant red light.

## Results

### Auxin is not sufficient to stimulate adventitious rooting in Norway spruce hypocotyl cuttings kept under white light

To investigate the role of light in adventitious root initiation (ARI) in Norway spruce, we used de-rooted Norway spruce seedlings as a model system. Three-week-old seedlings grown under white light condition were harvested and de-rooted as described in Materials and methods (Fig. S1). They were first kept in hormone free (HF) distilled water, under three different white light (WL) regimes as described in Materials and methods (irradiance 69 to 75 μM/m^2^/s^−1^; Supplementary Fig. S2A,B,C). In these conditions no AR developed at the base of the hypocotyls regardless of the photoperiod or the source of white light. Since we obtained the same results in all WL conditions, we continued to use constant WL (cWL) provided by LEDs (Supplementary Fig. S1D and Supplementary Fig. S2C) as control conditions for the rest of the study.

It is well known that auxin can stimulate adventitious rooting in several plant species (Lakehal & Bellini, 2019), therefore we tested whether different types of auxin could stimulate ARI under cWL. Three-week-old de-rooted seedlings were treated with 1 or 5 μM of Indole 3-Acetic Acid (IAA), 1-Naphthalene 3-Acetic Acid (1-NAA) or Indole 3-Butyric Acid (IBA), but none of the auxins could induce ARI under cWL conditions (Fig. 1A). These results indicate that auxin is not sufficient to induce ARI in de-rooted Norway spruce seedlings kept under cWL.

**Fig 1.**
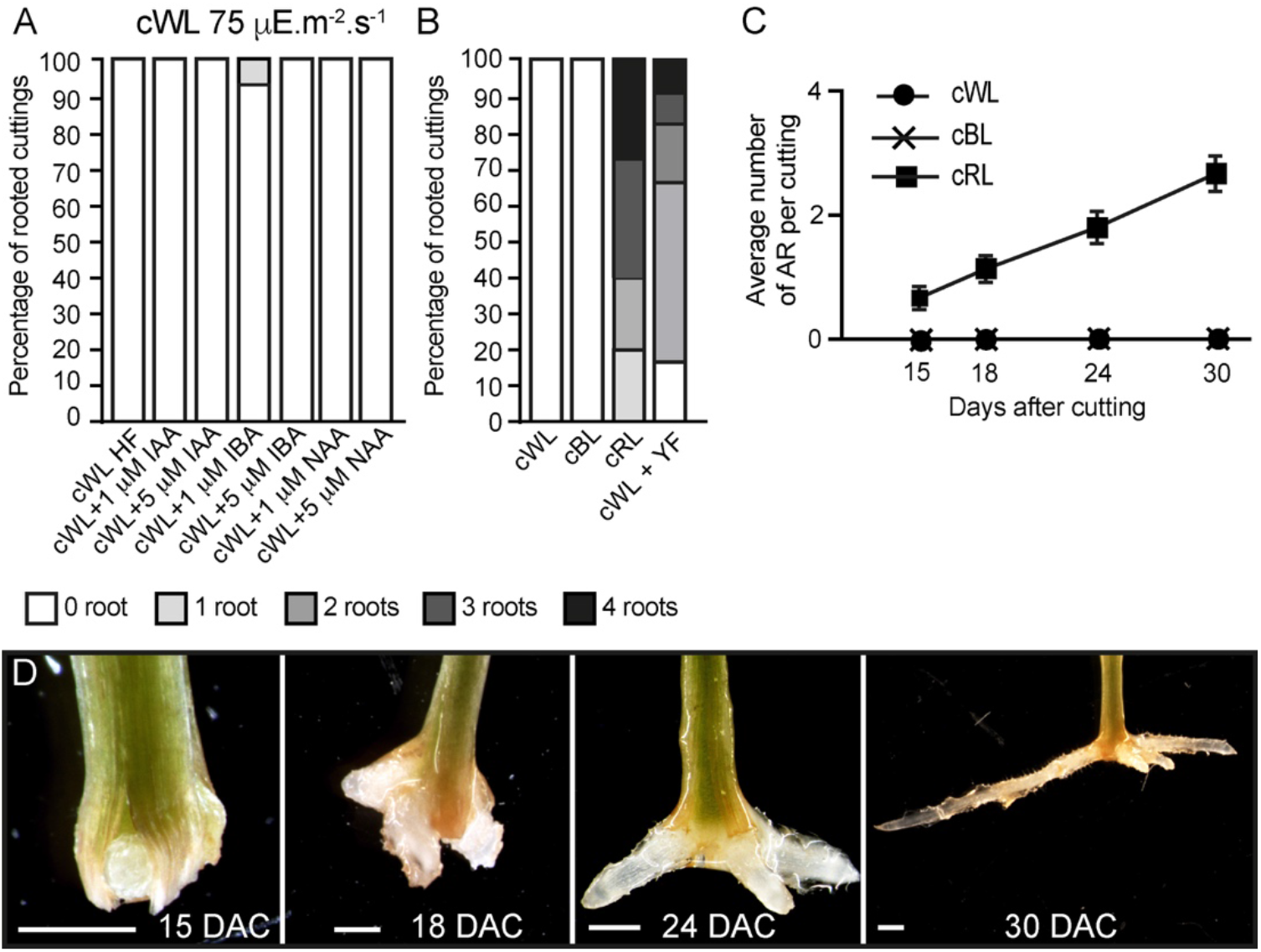
Constant blue light inhibits, while constant red light promotes, ARI in de-rooted Norway spruce hypocotyls. (A) Percentage of rooting in hypocotyl cuttings kept under constant white light (cWL). Three-week-old Norway spruce seedlings were de-rooted and kept for 30 days in hormone free (HF) distilled water, or in the presence of 1 or 5 μM indole-3-acetic acid (IAA), indole-3-butyric acid (IBA) or 1-naphthalene acetic acid (NAA), under cWL. (B) Percentage of rooting in hypocotyl cuttings kept under cWL, cWL with a yellow filter (YF), constant red light (cRL) or constant blue light (cBL). Three-week-old Norway spruce seedlings were de-rooted and kept for 30 days in hormone free (HF) distilled water under the different light conditions. (C) Average number of ARs on Norway spruce hypocotyls kept under cWL, cBL or cRL. Three-week-old Norway spruce seedlings were de-rooted and kept for 30 days in hormone free (HF) distilled water. The hypocotyl cuttings were observed every day to count emerging AR primordia and emerged ARs as in (**d**). Fifteen seedlings were analyzed for each condition. (D) Different stages of AR development in de-rooted 3-week-old Norway spruce seedling in distilled water, under cRL. DAC: days after cut; scale bar = 2 mm.

### Constant red light (cRL) and constant blue light (cBL) have opposite effects on ARI in de-rooted Norway spruce hypocotyls

Next, we wondered whether a particular region of the light spectrum had a negative effect on ARI. To test this hypothesis, three-week old de-rooted seedlings were kept in HF distilled water under either cWL (Fig. S1D and Fig. S2C), constant blue light (cBL) (Fig. S1f and Fig. S2D) or constant red light (cRL) (Fig. S1E and Fig. S2E). Interestingly, AR developed only when seedlings were kept under cRL conditions (Fig. 1B-D). In these conditions 100% of the cuttings developed at least 1 AR by 30 days after cutting (DAC) (Fig. 1B). These results raised the possibility that in cWL, the blue light part of the spectrum has a negative effect on ARI. To test this hypothesis, hypocotyl cuttings were kept under cWL but a yellow filter was added to remove the blue wavelength region (Fig. S2F). In these conditions 83% of the cuttings developed at least 1 AR (Fig. 1B).

These results indicate that cRL promotes ARI, whereas cBL inhibits this process, and the absence of AR under cWL could be due to the presence of blue light.

### Constant red light induces ARI by reducing endogenous levels of jasmonate, cytokinins, and abscisic acid

We have previously shown that ARI in intact etiolated Arabidopsis hypocotyls is under the control of IAA and JA signaling pathways (Gutierrez *et al*., 2012). More recently we showed that JA and CK cooperatively repress this process (Lakehal *et al.*, 2020a). We therefore hypothesized that light might also modulate phytohormone signaling and/or homeostasis pathways to control ARI in Norway spruce hypocotyls. To test this hypothesis, we quantified the endogenous contents of different hormones known to either inhibit ARI, such as JA, abscisic acid (ABA) or CKs (Steffens *et al*., 2006; Ramírez-Carvajal *et al*., 2009; Gutierrez *et al*., 2012; McAdam *et al*., 2016; Mao *et al*., 2019) or promote ARI, such as IAA or salicylic acid (SA) (Gutierrez *et al*., 2012). Quantifications were performed for hypocotyl samples taken at time 0 (T0), i.e. just before being transferred in HF distilled water, and at the base of hypocotyl cuttings maintained under different light qualities 6, 24, 48 and 72 hours after cutting (Fig. 2; Supplemental data sets 1 and 2).

**Fig 2.**
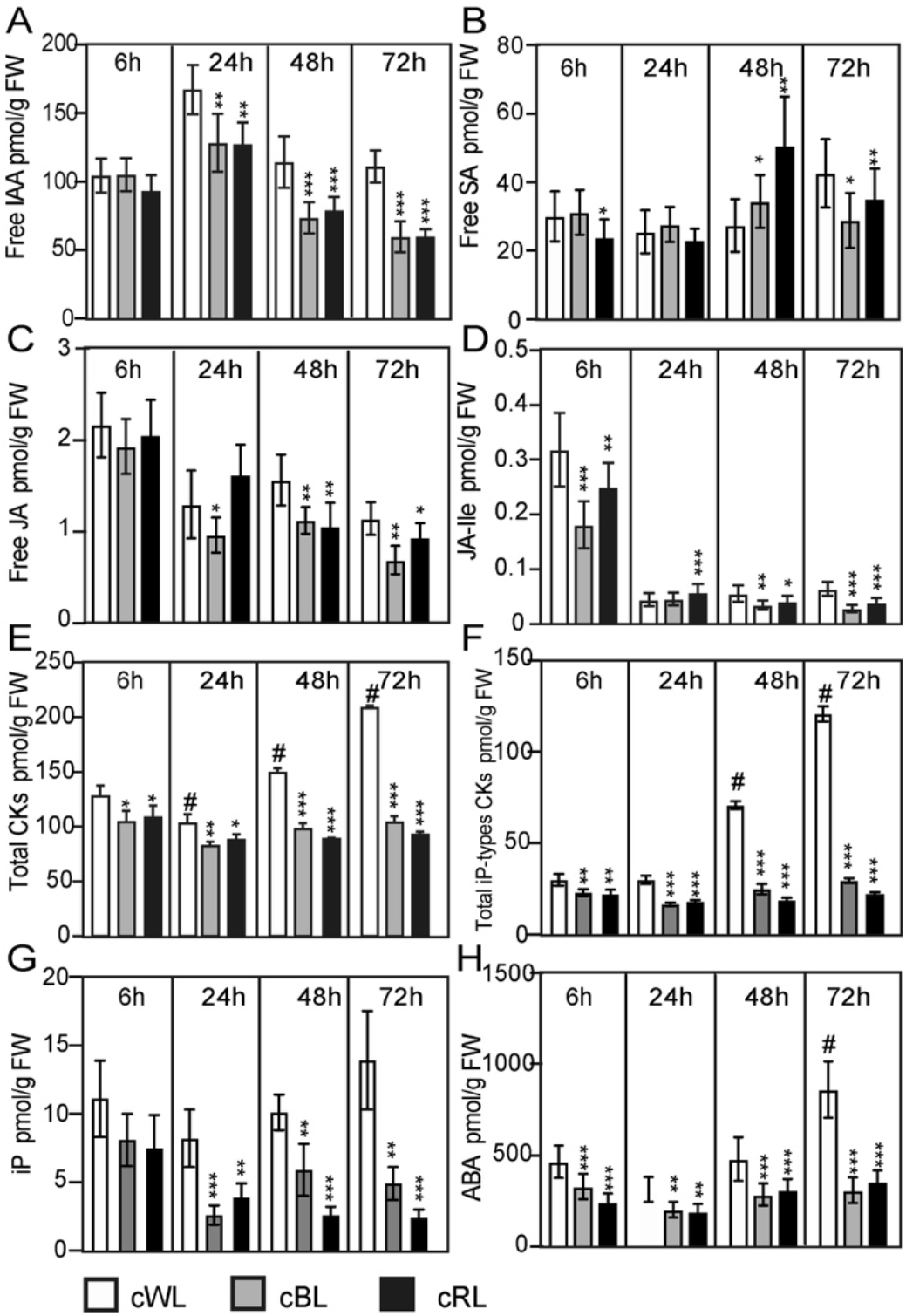
Light spectral quality controls the homeostasis of endogenous contents of different hormones, which positively or negatively affect ARI. (A) to (H) Three-week-old Norway spruce seedlings were grown under long day conditions as described in Materials and methods. De-rooted seedlings were then kept in constant cWL, cBL, or cRL for 6, 24, 48 and 72 hours. For each time point in each condition 5 mm hypocotyls were taken and pooled for hormone quantification. Five independent biological replicates were collected for analysis of IAA, ABA, SA and JA contents, and additional five biological replicates were collected for cytokinin quantification. (A) free IAA content; (B) free salicylic acid (SA) content; (C) free jasmonic acid (JA) content; (D) jasmonate-isoleucine (JA-Ile) content; (E) content of total cytokinins (CKs); (F) total isopentyl-adenine type (iP-type) CKs; (G) isopentyl-adenine (iP) content; (H) Free abscisic acid (ABA) content. Values are the means with standard deviation (SD) of 5 biological replicates. Asterisks indicate statistically significant differences under light conditions cRL versus cWL and cBL versus cWL in t-test; *, **, and *** correspond to P-values of (0.05 > p > 0.01, 0.01 > p > 0.001, and p < 0.001 respectively; *n* = 5); Dashes indicate statistically significant difference 6 h versus 24 h, 48 h or 72 h in t-test; ^#^, ^##^, ^###^ correspond to P-values of (0.05 > p > 0.01, 0.01 > p > 0.001, and p < 0.001, respectively; n = 5; FW, fresh weight.

Under cWL conditions, we observed a 40% increase in the free IAA level 6 h after cutting compared to the level at T0 (Supplemental data set 1), but thereafter the IAA content remained mostly constant over time, with the exception of a slight increase 24 h after cutting (Fig. 2A; Supplemental data set 1). There was no effect of wounding on the endogenous content of SA, as shown in Supplemental data set 1, and the SA content remained constant up to 48 h after cutting (Fig. 2B). A slight increase could be observed 72 h after cutting (Fig. 2B, Supplemental data set 1). Based on these results, we concluded that the inability to initiate ARs under cWL could not be explained by a reduction in auxin content, and this is in agreement with the fact that exogenous application of IAA or other auxins could not stimulate ARI under these conditions (Fig. 1A)

We have previously shown that JA and its bioactive form Jasmonoyol-Isoleucine (JA-Ile) inhibit ARI in Arabidopsis hypocotyls (Gutierrez *et al*., 2012; Lakehal *et al*., 2020a). Here, as expected in the case of wounding, JA and JA-Ile were induced 6 h after cutting in all light conditions (Supplemental data set 1). Under cWL conditions, we observed a 93 % increase in both JA and JA-Ile 6 h after cutting and transfer to cWL conditions (Supplemental data set 1). Both JA and JA-Ile contents subsequently decreased over time (Fig. 2C,D; Supplemental data set 1). Interestingly, although we observed a significant decrease (72%) in total CK content 6 h after cutting compared to the content at T0 (Supplemental data set 2), the total CK content then increased again and CKs accumulated over time at the base of the hypocotyls (Fig. 2). This increase in CKs was mostly due to the accumulation of isopentyl-adenine-type (iP-type) cytokinins including the precursors iP riboside 5’-monophosphate (iPRMP) and iP ribosides (iPR) (Fig. 2E-G; Supplemental data set 2). Similarly, we observed a 78 % decrease in the endogenous content of ABA 6 h after cutting compared to the content at T0 (Supplemental data set 2), but it increased again over time (Fig. 2h).

Interestingly, in cRL conditions, although the free IAA content was significantly reduced compared to that in cWL 24 h after cutting and continued to decrease over time (Fig. 2A), cutting developed ARs. This could be explained by a significant reduction in the amount of JA, Ja-Ile, CKs and ABA compared to cWL at all time points (Fig. 2C-H). In contrast to what was observed in cWL, the amounts of CKs and ABA remained constant over time, except for iP which continued to decrease over time (Fig. 2G). Although the interaction between light spectral quality and hormone homeostasis is complex, these results suggest that the positive effect of cRL on ARI cannot be explained by the modification of IAA homeostasis; rather it is a consequence of a decrease in negative regulators such as JA, JA-Ile, and iP.

Surprisingly, although cBL had the same effect as cRL on the endogenous levels of hormones (Fig. 2A-H), ARs could not develop. These results suggest that cBL inhibits adventitious rooting through another pathway yet to be identified. In the remainder of this study we therefore investigated only the role of cRL on ARI.

### Exogenously applied JA and CK inhibit ARI under cRL

To get physiological insights into the possible crosstalk between IAA, JA and CKs during cRL-induced AR development, we treated the de-rooted seedlings exogenously with either auxins, JA, or CK alone or in combination.

First, we showed that the three types of auxin (IAA, NAA and IBA) enhanced AR formation under cRL conditions in a dose-dependent manner (Fig. 3A-C). IBA, which is often used to induce rooting in recalcitrant species (reviewed in Geiss *et al*., 2018; Stevens *et al*., 2018), appeared to be the most efficient auxin (Fig. 3A). When applied exogenously, both JA and 6-Benzylaminopurine (6-BAP) inhibited AR development in a dose-dependent manner (Fig. 3D,E) and they both repressed the positive effect of exogenous IBA (Fig. 3F,G). These results suggest that JA and CK act downstream of auxin signaling to repress ARI, as has been described for intact Arabidopsis hypocotyls (Gutierrez *et al*., 2012; Lakehal *et al*., 2020a). When JA and CK were combined, no additive or synergistic effect was observed (Fig. 3H), suggesting that they act in the same pathway. These results are in agreement with our previous data that showed that CK signaling is induced by JA to repress AR formation in intact Arabidopsis hypocotyls (Lakehal *et al*., 2020a).

**Fig 3.**
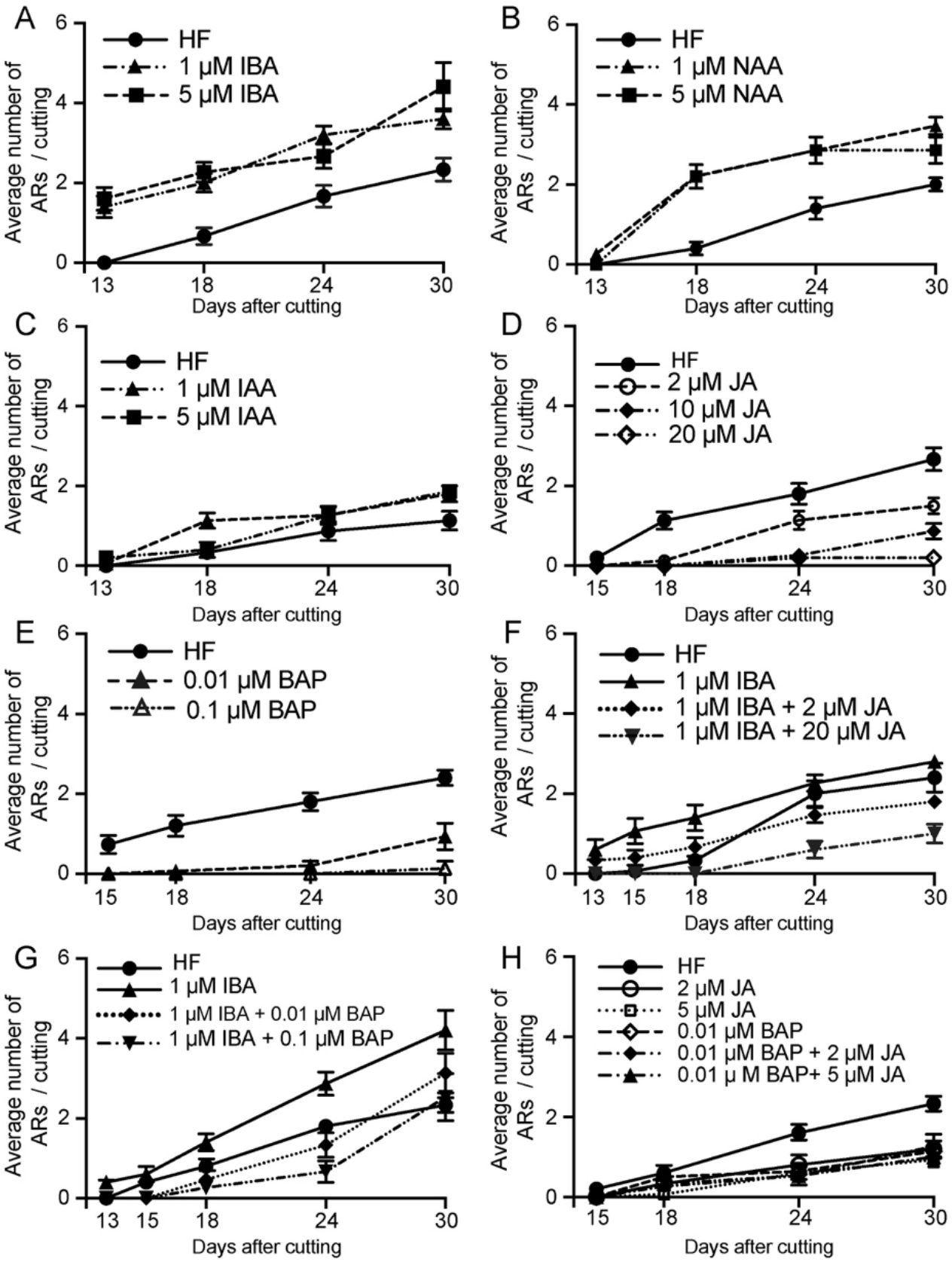
Auxin enhances RL-induced adventitious rooting, whereas CKs and JA repress this process. Three-week-old Norway spruce seedlings were de-rooted and kept under cRL for 30 days in hormone free (HF) distilled water, or in the presence of 1 or 5 μM IBA (A); or in the presence of 1 or 5 μM NAA (B); or in the presence of 1 or 5 μM IAA (C); or in the presence of jasmonate (JA) (D); or in the presence of benzyl adenine (BAP) (E); or in the presence of both JA and IBA (F); or in the presence of both BAP and IBA (G); or in the presence of both JA and BAP (H). (A) Both concentrations of exogenously applied IBA significantly increased the average number of ARs over time (repeated measures ANOVA; P < 0.0001). (B) Both concentration of exogenously applied NAA significantly increased the average number of adventitious roots over time (repeated measures ANOVA; P < 0.0001). (C) Exogenously applied IAA did not have any significant effect compared to HF conditions (Repeated measures ANOVA; P = 0.073). (D) Exogenously applied JA had a significant negative effect on the average number of ARs over time (repeated measures ANOVA; P < 0.0001). This negative effect was concentration dependent (repeated measures ANOVA; P < 0.0001). (E) Exogenously applied BAP significantly reduced the number of ARs compared to HF conditions (repeated measures ANOVA; P < 0.0001). (F) When JA was applied together with IBA, it significantly repressed the positive effect of IBA. The combination 1 μM IBA + 2 μM JA significantly reduced the number of roots compared to those in HF conditions by 24 days after cut (DAC) and 30 DAC (t-test, degree of freedom df = 14; p-value = 0.02 and 0.001 respectively). The combination 1 μM IBA + 20 μM JA significantly reduced the average AR number compared to HF conditions (t-test, df = 14; p-value < 0.01). (G) When BAP was applied together with IBA, it repressed the positive effect of IBA. The combination 1 μM IBA + 0.01 μM BAP significantly reduced the average number of ARs compared to the 1 μM IBA treatment. (13, 15, 18, 24 and 30 DAC; t-test, df = 14; p-value < 0.009, 0.007, 0.003, 0.0001And 0.02 respectively). The combination 1 μM IBA + 0.1 μM BAP significantly inhibited rooting induced by 1μM IBA at (13,15,18,24 and 30 DAC (t-test, df = 14; p-value < 0.008,0.007,0.0009, 0.000000009 and 0.0009 respectively); (H) Both JA and BA repressed AR compared to the HF control, but no significant additive or synergistic effect was observed when JA and BAP were applied together. The combination 0.01 μM BAP + 2 μM JA significantly inhibited rooting compared to the HF control at 24 and 30 DAC (t-test, df = 14; p-value < 0.0007 and 0.003 respectively), but the result was not significantly different from JA or BAP alone; The combination of 0.01 μM BAP + 5μM JA significantly inhibited rooting compared to the HF controls at 24 and 30 DAC (t-test, df = 14; p-value < 0.00004 and 0.0000003 respectively), but with no significant difference from JA or BAP alone. (A to H) Results are expressed as average number of ARs per cutting. Error bars indicate standard error (SE). For each treatment 15 hypocotyls were used, and the experiment was repeated twice.

### JA signaling is downregulated in de-rooted Norway spruce hypocotyls kept under cRL compared to cWL conditions

The reduction in content of JA and JA-Ile under cRL compared to cWL (Fig. 2B) prompted us to check whether the expression of genes involved in JA biosynthesis or JA signaling was affected in hypocotyls of seedlings kept under cRL compared to cWL conditions. We first searched the Norway spruce genome (Nystedt *et al*., 2013) for putative orthologs of the key Arabidopsis genes in JA signaling or biosynthesis using http://congenie.org/. We looked for putative orthologs of the JA-Ile receptor *CORONATINE INSENSITIVE 1 (AtCOI1)* (Xie *et al*., 1998)*, AtMYC2* transcription factors (Lorenzo *et al*., 2004), the *JASMONATE ZIM-DOMAIN3 (AtJAZ3) and AtJAZ10* transcriptional repressors (Chini *et al.*, 2007; Thines *et al.*, 2007; Yan *et al.*, 2007) and *ALLENE OXYDE CYCLASE (AtAOC)*, which encodes a key enzyme in the JA biosynthesis pathway (Stenzel *et al*., 2012). We found that the Norway spruce genome contains eleven putative *PaCOI1-like* genes (Fig. S3A), five putative *PaMYC2-like* genes (Fig. S3B), one and three putative orthologs of *PaJAZ3-like* and *PaJAZ10-like* respectively (Fig. S3C) and two putative *PaAOC-like genes* (Fig. S3D).

The full-length coding sequences are reported in Fig. S4. When more than one putative ortholog was identified we chose that most closely related to the Arabidopsis one for further experiments.

We first confirmed that expression of *PaMYC2-like* (MA_15962g0010), *PaJAZ3-like* (MA_6326g0010), *PaJAZ10-like* (MA_10229741g0010) and *PaAOC-like* (MA_56386g0010) was induced by exogenously applied JA (Fig. 4A). We next checked the expression of these genes, together with *PaCOI1-like*, at the base of hypocotyl cuttings kept under cWL or cRL for 6 h, 24 h, 48 h or 72 h after cutting (Fig. 4B). The relative amounts of *PaMYC2-like, PaJAZ3-like*, and *PaAOC-like* transcripts were slightly reduced in cRL compared to those observed in cWL (Fig. 4B). These results are in agreement with the reduced content of JA-Ile, the active form of JA, in cRL compared to cWL (23% less in cRL compared to cWL) (Fig. 2F and Supplemental data set 2). Twenty-four hours after cutting, *PaMYC2-like, PaJAZ10-like*, and *PaCOI1-like* were slightly upregulated in cRL compared to cWL (Fig. 4b), results which also coincided with a slightly higher concentration of JA-Ile (20% more in cRL compare to cWL) (Fig. 2F). Later on, by 48h and 72h, the JA responsive genes were downregulated (Fig. 4), and this coincided with a greater decrease in JA and JA-Ile contents in cRL compared to cWL (Fig. 2F; Supplemental data set 2). In conclusion, when de-rooted hypocotyls were kept in cRL conditions, the endogenous contents of JA and JA-Ile decreased faster than in cWL, resulting in downregulation of JA signaling in cRL compared to cWL conditions. According to our previous results (Gutierrez *et al*., 2012; Lakehal *et al*., 2020a), this most likely contributes to an improvement in adventitious root development.

**Fig 4.**
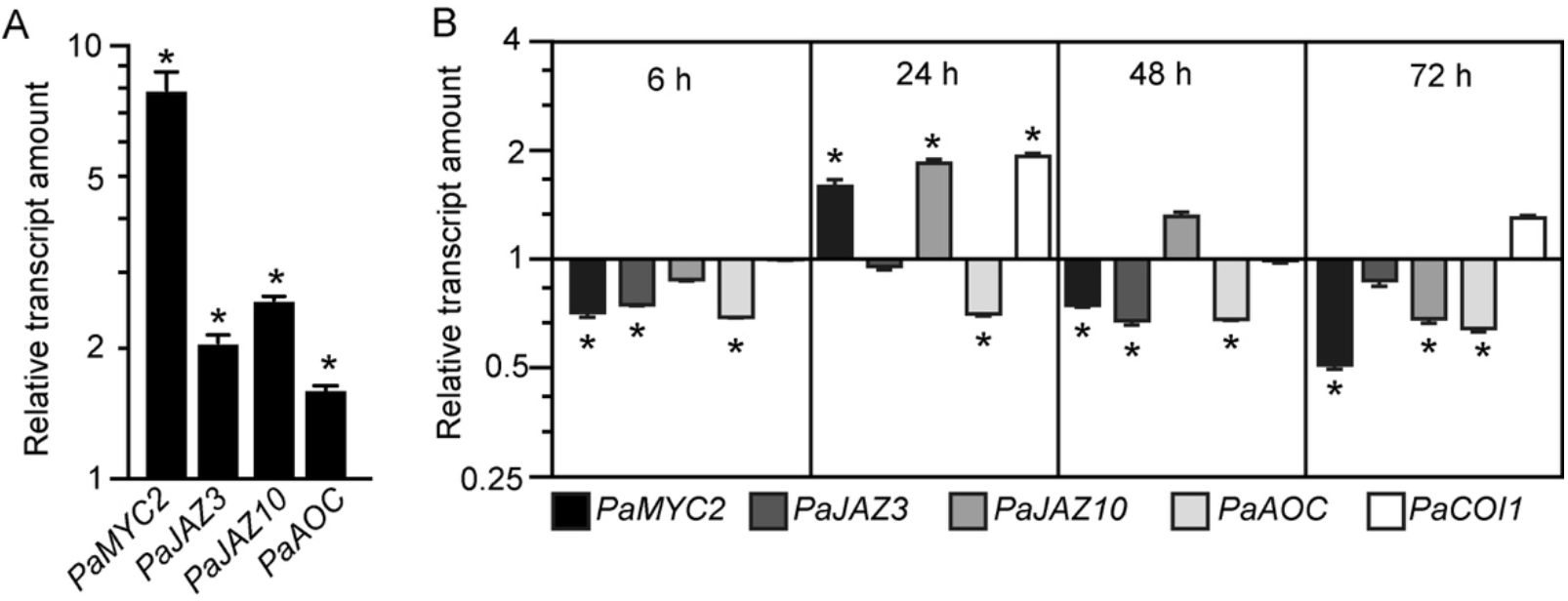
JA signaling is downregulated in hypocotyl cuttings kept under constant red light (cRL) compared to hypocotyl cuttings kept under constant white light (cWL) (A) Relative transcript amount of putative JA-responsive genes *PaMYC2-like, PaJAZ3-like, PaJAZ10-like* and *PaAOC-like* in three-week-old Norway spruce seedling hypocotyls treated for 3 h with 50 μM JA or with a mock treatment. Values are relative (on a log10 scale) to the values for mock treated seedlings which were arbitrarily set to 1. Error bars indicate ±SE obtained from two independent biological replicates, with three technical replicates of each. A *t*-test was carried out, and values indicated by asterisks differed significantly from the mock values (P < 0.001, n ≥ 80). (B) Relative transcript amounts of putative JA-responsive genes *PaMYC2-like, PaJAZ3-like, PaJAZ10-like* and *PaAOC-like*, and the putative JA receptor *PaCOI1-like*, at the base of three-week-old de-rooted hypocotyls kept under cRL or cWL for 6, 24, 48 and 72 h. The values are relative (on a log2 scale) to the values obtained from seedlings kept under cWL, which were arbitrarily set to 1. Error bars indicate ±SE obtained from two independent biological replicates, with three technical replicates of each. A *t*-test was carried out, and asterisks indicate values that differed significantly from the control values (P < 0.001, n ≥ 80).

### Exogenously applied JA inhibits ARI in Norway spruce hypocotyl in cRL

Our data indicated that JA signaling was rapidly downregulated (within 6 hours) in the hypocotyls of cuttings kept under cRL compared to cWL, suggesting that JA signaling repressed an early event in AR formation. To assess the repressive effect of JA at the cellular level, we analyzed and compared the anatomy of hypocotyls kept under cRL, in hormone free distilled water or in the presence of 1 μM IBA or 20 μM JA, at different time points after cutting (Fig. 5 and 6). On day 0 (i.e. immediately after cutting the hypocotyl of three-week-old Norway spruce seedlings), the hypocotyl had a primary structure consisting of a single layer of epidermis, and six to seven cortex cell layers composed of isodiametric parenchyma cells (Fig. 5A). We could guess at the presence of the endodermis layer although it was not always morphologically distinctive (Fig. 5A). The central cylinder consisted of seven to eight layers of parenchyma cells which were smaller compared to the cortex parenchyma cells. The vascular system was composed of a continuous procambium and differentiated elements of the protophloem and protoxylem. The pith region showed small isodiametric parenchyma cells (Fig. 5A).

**Fig 5.**
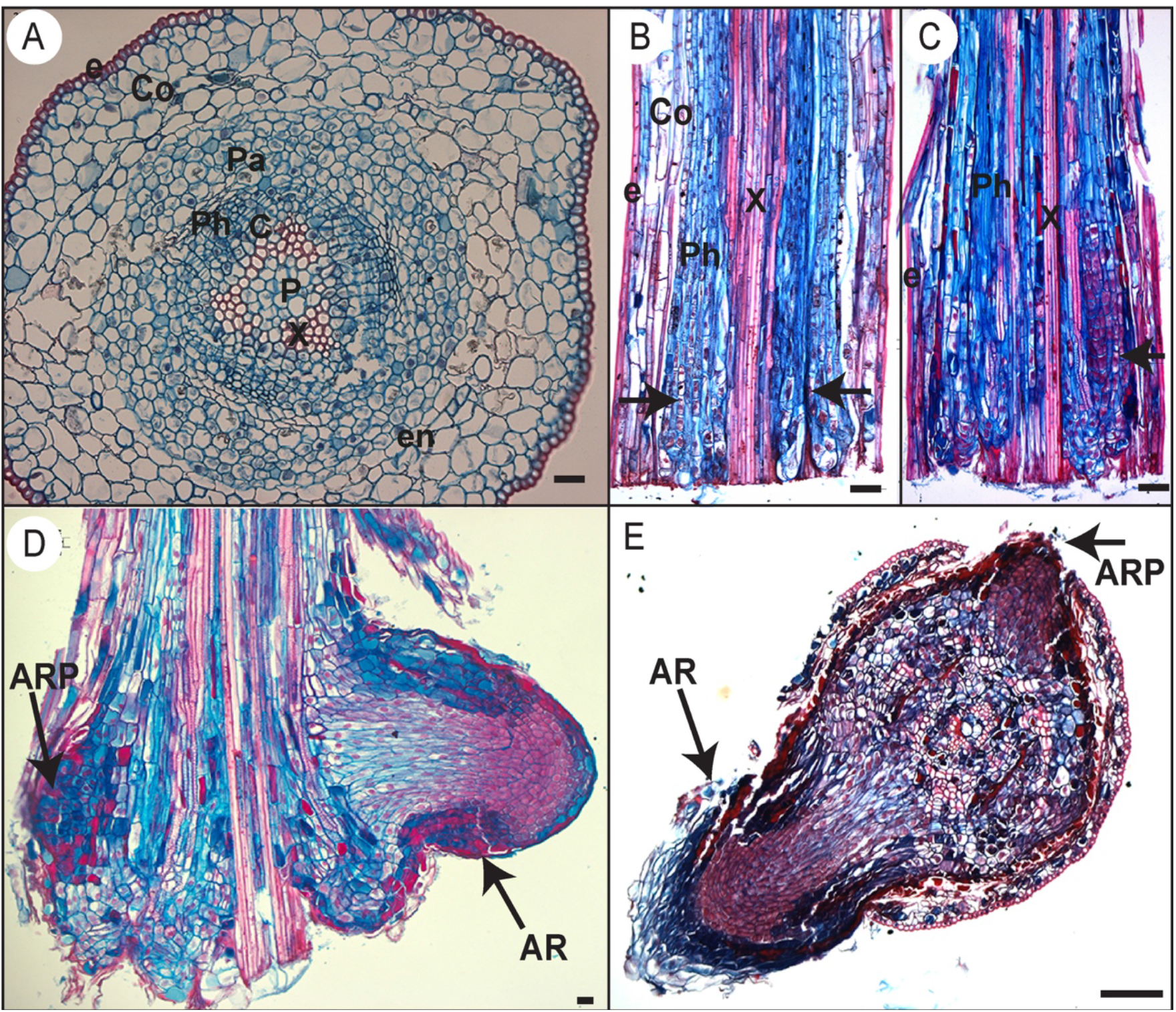
Histological analysis of AR development in Norway spruce hypocotyl cuttings under cRL. (A) Cross section through the base of a 21-day-old hypocotyl cutting just before transfer to rooting conditions. (B) Longitudinal section through the base of a hypocotyl cutting kept in hormone free (HF) conditions for 5 days. Arrowheads indicate localized cell divisions, probably at the points of origin of future ARs. (C) Longitudinal section through the base of a hypocotyl cutting kept in the presence of 1 μM IBA for 10 days. Arrowheads indicate the presence of meristemoids. (D) Longitudinal section through the base of a hypocotyl cutting kept in the presence of 1 μM IBA for 13 days. Arrows indicate an adventitious root primordium (ARP) and an emerged adventitious root (AR). (E) Cross section through the base of a hypocotyl cutting kept in HF conditions for 15 days. Arrows indicate an ARP and an emerged AR. e, epidermis; Co, cortex; Pa, parenchyma; Ph, phloem; en, endodermis; c, cambium region; P, pith; x, xylem. In all panels scale bars = 20 μm.

Three days after cutting, in HF distilled water and in the presence of 1 μM IBA, a few cells in the cambial zone and in the adjacent phloem cells developed dense cytoplasm and large nuclei indicating the initiation of the first cell divisions (Fig. 6A,B). In contrast, no obvious histological modifications were observed in the presence of 20 μM JA, and the histological organization remained unchanged compared to day 0 (Fig. 6C). By five days after cutting, periclinal and anticlinal divisions were clearly observed in the cambial zone and in the outermost layers of the phloem region (Fig. 6D). In longitudinal section these cells were seen to be organized into vertical files external, but adjacent, to the vascular cylinder (Fig. 5B). In the presence of 1 μM IBA it was clearly apparent that the number of cell divisions was higher than under HF conditions (Fig. 6E), whereas no divisions were observed in the presence of 20 μM JA (Fig. 6F). Ten days after cutting, in HF conditions, cambial derivatives divided and formed radial rows of tracheal elements around the xylem, but no AR meristemoids were observed (Fig. 6G), while in the presence of 1 μM IBA several AR root meristemoids were visible (Fig. 5C and Fig. 6H); no cell divisions were observed in the presence of 20 μM JA (Fig. 6I). It was only by thirteen days after cutting that AR meristemoids began to form at the periphery of the trachea elements in HF conditions (Fig. 6J), while in the presence of IBA, AR had already emerged (Fig. 5D and Fig. 6K); still no cell division activity was observed in the presence of JA (Fig. 6L). In HF, the first emergence of ARs was observed only 15 days after cutting (Fig. 5E). These data clearly indicate that exogenously applied JA repressed the very early stage of ARI, whereas auxin not only promoted but also speeded up the process.

**Fig 6.**
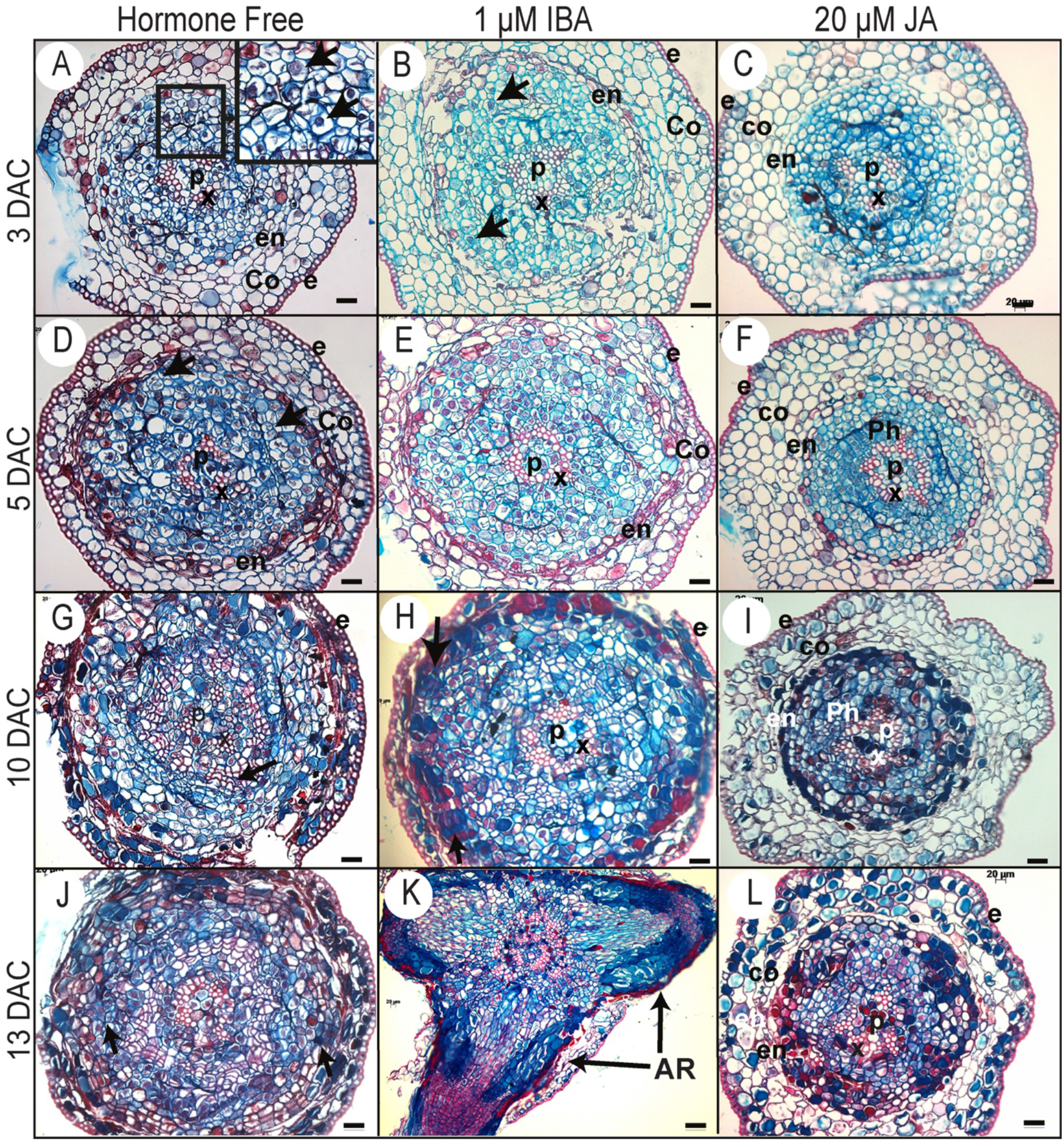
Histological analysis of ARI in Norway spruce hypocotyl cuttings under cRL in the presence of IBA, JA or HF. (A, D, G, J) Cross sections were prepared from the base of de-rooted Norway spruce hypocotyl cuttings, kept under cRL in hormone free (HF) distilled water for 3, 5, 10 or 13 days. In (A) the arrows show early cell activation, *i.e.* small cells with dense cytoplasm and large nuclei. In (D) arrows indicate anticlinal and periclinal division. In (G) arrows show the presence of tracheal elements. In (j) arrows indicate meristemoids developing at the periphery of the tracheal elements. (B, E, H, K) Cross sections were taken from the base of de-rooted Norway spruce hypocotyl cuttings, kept under cRL in the presence of 1 μM IBA for 3, 5, 10 or 13 days. In (B) arrows indicate cell divisions; In (H) arrows indicate the presence of meristemoids. In (K) arrows indicate emerged adventitious roots (ARs). (C, F, I, L) Cross sections were prepared from the base of de-rooted Norway spruce hypocotyl cuttings, kept under cRL in the presence of 20 μM JA for 3, 5, 10 or 13 days. e, epidermis; Co, cortex; Ph, phloem; en, endodermis; c, cambium region; P, pith; x, xylem. In all panels scale bars = 20 μm.

## Discussion

When optimizing conditions for the development of roots from cuttings during vegetative propagation, it has long been considered that light should be taken into account as an important factor. Since the early eighties, studies related to the effect of light on AR formation in woody plant cuttings have been pursued (Druart *et al*., 1982; Hammerschlag, 1987; Klopotek *et al*., 2010). Several studies have addressed the effect of light quality and/or intensity on the rooting of cuttings (Jarvis & Shaheed, 1987; Baraldi *et al*., 1988; Fuernkranz *et al*., 1990; Fett-Neto *et al*., 2001; Iacona & Muleo, 2010; Daud *et al*., 2013), but the exact molecular and genetic mechanisms underlying the role of light as well as the downstream targets of light signals during ARI remain poorly understood. In this study, we showed that under cWL conditions de-rooted Norway spruce hypocotyls were unable to produce AR and that this was unlikely to be due solely to an increase in endogenous CK levels, as suggested by (Strömquist & Eliasson, 1979; Bollmark & Eliasson, 1990) since exogenously applied auxin was unable to stimulate AR development. This result led us to hypothesize that specific wavelengths within the white spectrum could be responsible for this strong inhibition, and we therefore grew seedlings under white or monochromatic LEDs. In these conditions, we showed that AR developed under cRL even in the absence of exogenous auxin application, while they did not develop under cBL.

The cRL-mediated ARI is possibly due to a faster reduction in JA and JA-Ile contents compared to cWL. cRL also repressed the accumulation of CKs and ABA which was observed under cWL. Reduction in the amounts of the three negative regulators to below a certain threshold led to a de-repression of ARI.

The reduced amount of JA and JA-Ile in the base of cuttings could be partially explained by downregulation of *de novo* JA biosynthesis, since we observed a reduced amount of the JA precursor *cis*-12-oxo-phytodienoic acid (*cis*-OPDA) under cRL compared to cWL conditions 24, 48 and 72 h after cutting (Supplemental data set **2**). This result is supported by the downregulation of expression of *PaAOC-like*, a key gene in the JA biosynthesis pathway. The link between light and the JA signaling pathway has been proposed in several contexts, but how it affects JA biosynthesis is not yet known. Interestingly, several reports have revealed mechanistic insights into the roles of RL or red/far-red light ratio in the control of plant development or in adjusting growth-defense tradeoffs (Kazan & Manners, 2011). For example, (Campos *et al*., 2016) found that mutating the RL-receptor *Phytochrome B (PhyB)* suppressed the stunted rosette phenotype of the *JazQ* mutant, which harbors five null mutations in JAZ family members. These data suggest that *PhyB-mediated* light signaling interacts with JA signaling to control growth in Arabidopsis. Under our conditions, we found that the expression levels of *PaMYC2-like*, a master transcriptional regulator of JA signaling, were slightly but significantly downregulated under cRL compared to cWL. This result could be explained either by a direct effect of RL on transcriptional regulation or by the reduced amount of the bioactive form JA-Ile present under these conditions. The latter assumption is supported by the fact that exogenous application of JA strongly induced the expression of *PaMYC2-like*, an observation consistent with what has been reported in Arabidopsis. The reduced amounts of JA and JA-Ile and the downregulation of the expression of *MYC2-like* under cRL conditions led to a reduced JA response, as indicated by downregulation of the steady state level of expression of the well-known JA responsive genes (*PaJAZ10-like* and *PaJAZ3-like*).

Although the negative role of JA on ARI has been demonstrated in the model plant Arabidopsis, its exact role in other species previously remained unclear (reviewed in (Lakehal *et al*., 2020b). Our results showed that JA is a negative regulator of ARI in de-rooted Norway spruce hypocotyl and that it acts during the very early stages of initiation, as indicated by anatomical characterization. We showed that under cRL, certain cells, most likely cambial cells, have the competence to form ARs in a similar way to that described for other conifers such as *Pinus taeda, P. contorta* and *P. radiata* (Diaz-Sala *et al*., 1996; Lindroth *et al*., 2001B,a; Ricci *et al*., 2008). When we exposed the de-rooted hypocotyls to exogenous auxin, these cells exhibited rapid cell division and re-orientation of division planes to organize the AR meristem; in contrast when exogenous JA was added, no cell divisions were observed. In addition, we showed that JA repressed the positive effect of auxin when the compounds were applied together, supporting the conclusion that, as in Arabidopsis, JA acts downstream of auxin in the control of ARI in de-rooted Norway spruce hypocotyls. Moreover, we showed that, like JA, CK inhibits ARI and counteracts the action of auxin, but when combined with JA no additive or synergistic effect was observed, which is in agreement with our recent results showing that cytokinins act downstream of JA to repress ARI in Arabidopsis hypocotyls (Lakehal *et al*., 2020a).

Our data also showed that total CKs accumulated at the base of cuttings when the de-rooted seedlings were exposed to cWL, whereas their amount remained constant over time under cRL conditions. Under cWL, the increasing amounts of the iP-type CKs could be due to *de novo* biosynthesis since we also observed an increased level of the precursors iPR and iPRMP over time. (Bollmark & Eliasson, 1990) showed that de-rooted Norway spruce hypocotyls kept under high irradiance WL (270 μE.m^−2^.s^−1^) could not develop ARs, probably because of the accumulation of CKs. Nevertheless, the effect of WL may be more complex, since under our conditions, although the WL irradiance was much lower (75 μE.m^−2^.s^−1^), CKs still accumulated. In our conditions, as a response to wounding, JA and JA-Ile accumulated at the base of hypocotyls 6 h after cutting, and their content started decreasing thereafter, whereas iP-type CKs began to accumulate 48 hours after cutting. Recently we showed that MYC2-mediated JA signaling promotes the accumulation of CKs through control of expression of the *APETALA2/ETHYLENE RESPONSE FACTOR 115* transcription factor in Arabidopsis hypocotyls (Lakehal *et al*., 2020a). We therefore propose that in Norway spruce hypocotyls, wound induced accumulation of JA and JA-Ile triggers *de novo* cytokinin biosynthesis which results in inhibition of rooting. Interestingly, under constant cRL, the more rapid reduction in JA and JA-Ile contents and subsequent downregulation of MYC2-mediated JA signaling may prevent the induction of *de novo* CK biosynthesis and this would allow ARI, although we also cannot rule out a direct effect of cRL on the repression of CK accumulation.

In conclusion, we propose that de-rooted Norway spruce seedlings do not develop AR under WL, on the one hand because they accumulate CKs and ABA which are repressors of ARI and on the other because of the presence of light in the blue region of the spectrum which has a negative effect independently of JA, CKs and ABA. Understanding the role of blue light requires further investigation. In contrast, we showed that red light has a positive effect on ARI, most likely because it represses the JA signaling pathway and the accumulation of CKs.

## Supporting information

Supplemental data set 1

Supplemental data set 2

## Supplemental data

The following supplemental data are available

**Supplementary Fig. S1:** Method for *de novo* root organogenesis under different light conditions

**Supplementary Fig. S2:** Spectral emission curves for the various light sources used in this study

**Supplementary Fig. S3:** Phylogenetic trees

**Supplementary Fig. S4:** Coding sequences used for the phylogenetic analysis

**Supplementary Table S1** Primers used for qPCR experiments in this study

## Accession numbers

*AtMYC2* (At1g32640), *AtJAZ3* (At3g17860), *AtJAZ10* (At5g13220), *AtAOC1* (At3g25760), *AtCOI1* (At2g39940), *PaMYC2-like* (MA_15962g0010), *PaJAZ3-like* (MA_6326g0010), *PaJAZ10-like* (MA_10229741g0010) and *PaAOC-like* (MA_56386g0010)

## Acknowledgements

The authors sincerely thank Marta Derba Maceluch from the UPSC Microscopy platform for her advice and help with histological characterization.

This research was supported by the Ministry of Higher Education and Scientific Research in Iraq (to SA), by grants from the Swedish research councils FORMAS, VR, VINNOVA, Kempestiftelserna, and the Carl Tryggers Stiftelse (to CB), and by grants from the Ministry of Education, Youth and Sports of the Czech Republic (European Regional Development Fund-Project “Plants as a tool for sustainable global development” No. CZ.02.1.01/0.0/0.0/16_019/0000827), and the Czech Science Foundation (Project No. 19-00973S) (to ON)

