## Supplementary information for "Red light controls adventitious root regeneration by modulating hormone homeostasis in *Picea abies* seedlings"

**The following Supporting Information is available in this PDF:**

**Supplemental Fig. S1:** Method for *de novo* root organogenesis under different light conditions

**Supplemental Fig. S2:** Spectral emission curves for the various light sources used in this study

**Supplemental Fig. S3:** Phylogenetic trees

**Supplemental Fig. S4:** Coding sequences used for the phylogenetic analysis

**Supplemental Table S1** Primers used for qPCR experiments in this study

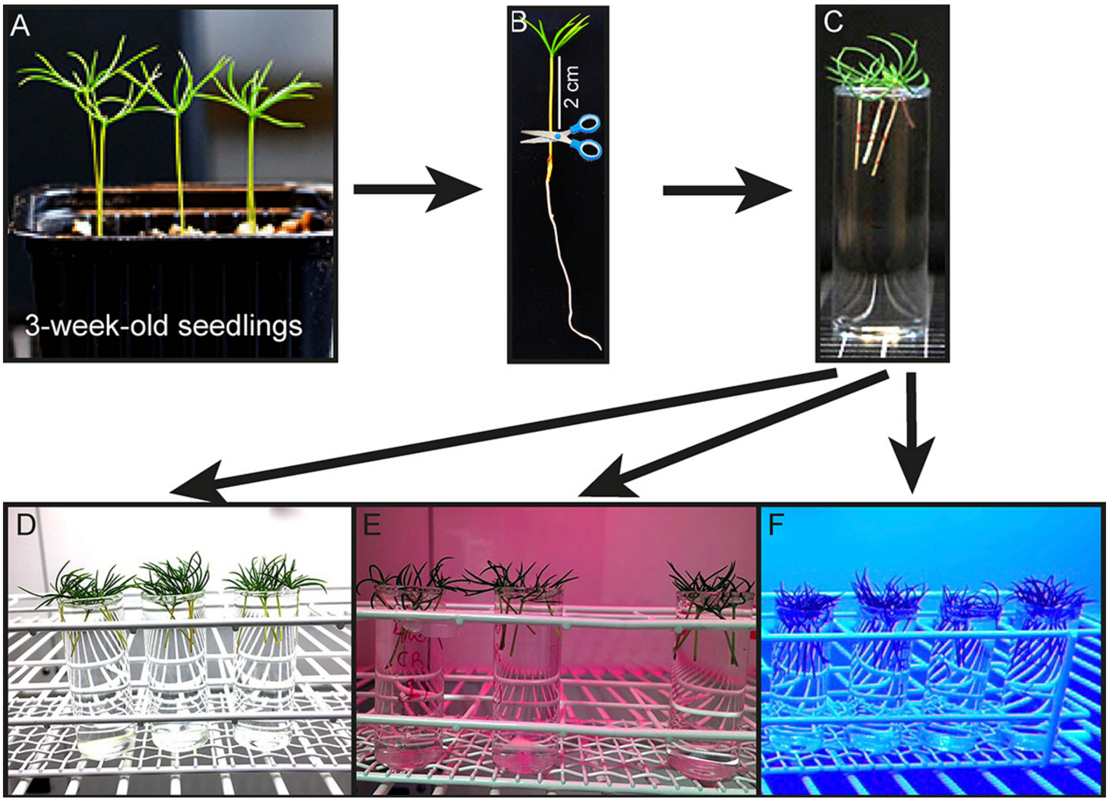


**Supplementary Fig. S1 Method for *de novo* root organogenesis under different light conditions**

(A) Three-week-old Norway spruce seedlings grown in vermiculite in a growth chamber under long day conditions

(B) Three-week-old Norway spruce seedling before cutting.

(C) Hypocotyl cuttings transferred to 24 ml vials filled with distilled water and placed in monochromatic cabinets equipped with white (d), red (e) or blue (f) LEDs

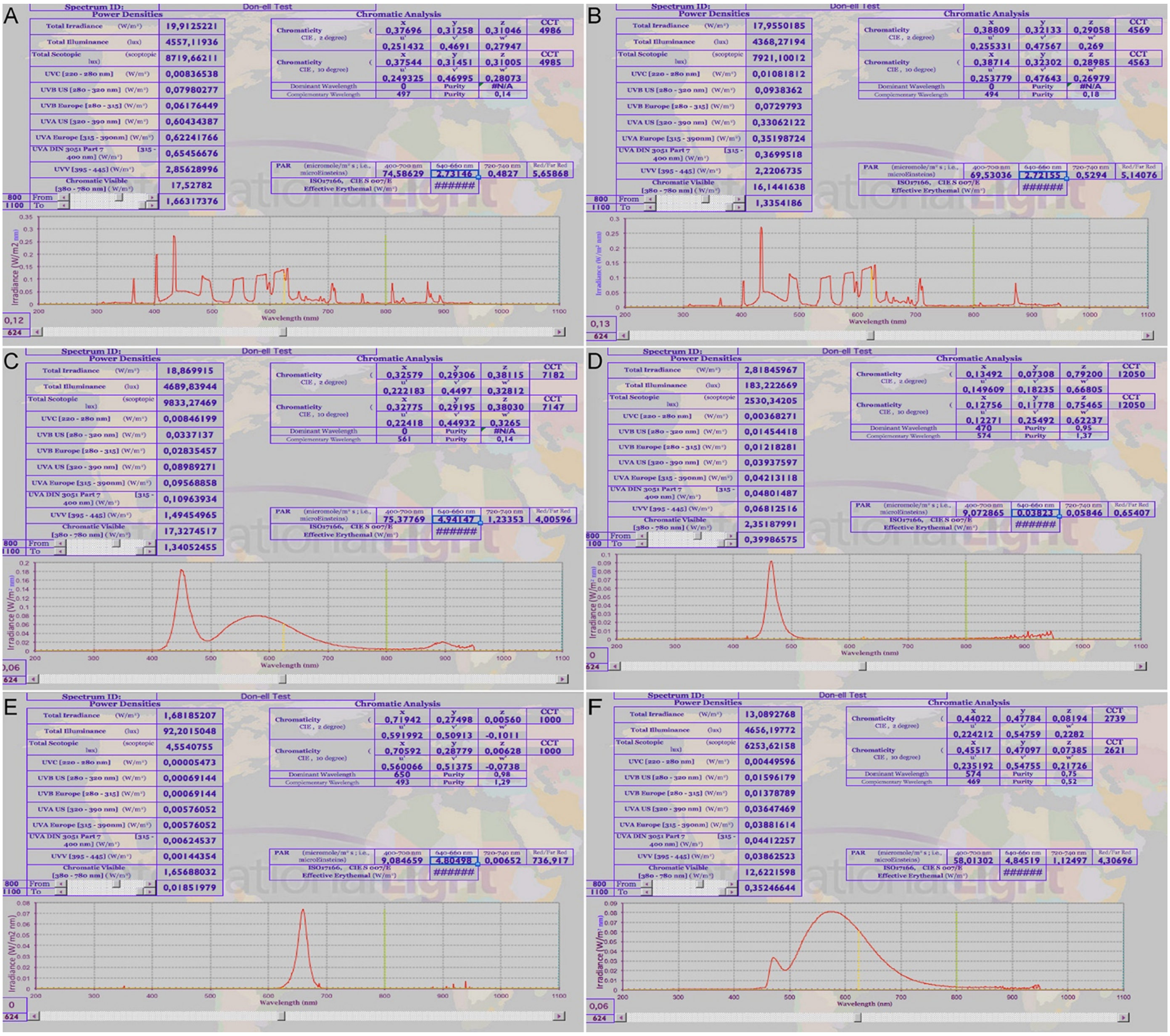

**Supplementary Fig. S2: Spectral emission curves for the various light sources used in this study.**

(A) White light TL-D 36W/840 Philips master, long days

(B) Cool white light F17T8/TL741, 17 W Philips, long days

(C) Continuous white light LED (cWL)

(D) Continuous white light LED + yellow filter

(E) Continuous blue light LED (cBL)

(F) Continuous red light LED (cRL).

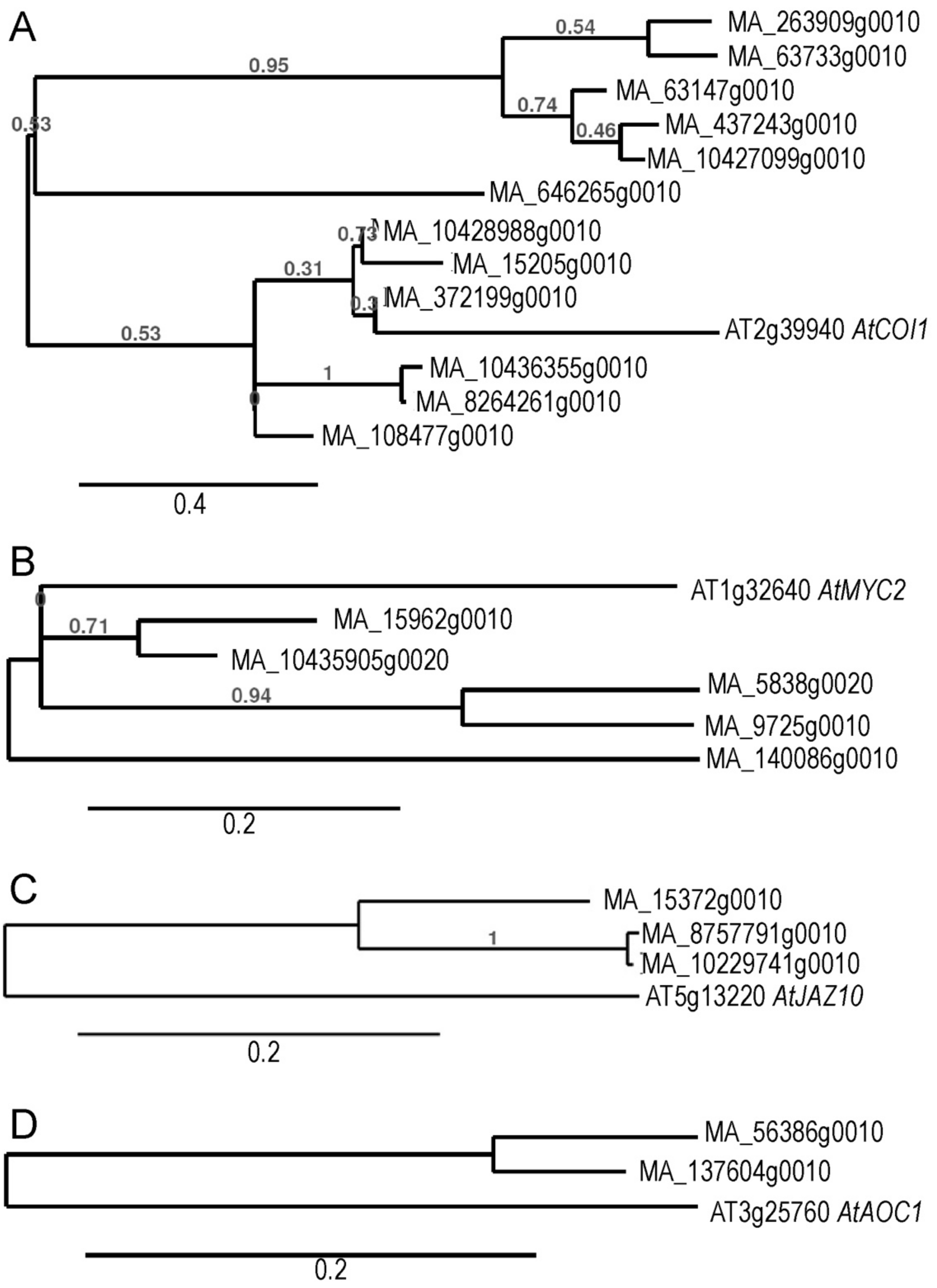


**Supplementary Fig. S3 Phylogenetic trees**

Neighbor-Joining phylogenetic trees (1000 bootstrap replicates), based on alignment of coding sequences of (A) *COI1* (B) *MYC2* (C) *JAZ10* and (D) *ACO1* (E) from *Picea abies* (MA) and *A. thaliana* (AT). The gene orthologs were named according to the http://congenie.org/ database. The sequences are provided in Supplemental Fig. S4.

**Supplementary Fig. S4 Coding sequences used for the phylogenetic analysis**

*Arabidopsis thaliana* and *Picea abies* coding sequences (CDS) were obtained from the TAIR (https://www.arabidopsis.org) and congenie (http://congenie.org/) databases respectively. To find putative Arabidopsis orthologs in *Picea abies*, a BLAST alignment of the coding sequences of selected genes was generated using "Genome Tools" at http://congenie.org/ with default settings. Subsequently, http://www.phylogeny.fr was used to construct phylogenetic trees based on the neighbor-joining method (with 1000 bootstrap replicates). Gene orthologs were named according to the http://congenie.org database.

**> COI1_ AT2G39940**

ATGGAGGATCCTGATATCAAGAGGTGTAAATTGAGCTGCGTCGCGACGGTTGATGATGTCAT
CGAGCAAGTCATGACCTATATAACTGACCCGAAAGATCGCGATTCGGCTTCTTTGGTGTGTC
GGAGATGGTTCAAGATTGATTCCGAGACGAGAGAGCATGTGACTATGGCGCTTTGCTACACT
GCGACGCCTGATCGTCTTAGCCGTCGATTCCCGAACTTGAGGTCGCTCAAGCTTAAAGGCAA
GCCTAGAGCAGCTATGTTTAATCTGATCCCTGAGAACTGGGGAGGTTATGTTACTCCTTGGG
TTACTGAGATTTCTAACAACCTTAGGCAGCTCAAATCGGTGCACTTCCGACGGATGATTGTC
AGTGACTTAGATCTAGATCGTTTAGCTAAAGCTAGAGCAGATGATCTTGAGACTTTGAAGCT
AGACAAGTGTTCTGGTTTTACTACTGATGGACTTTTGAGCATCGTTACACACTGCAGGAAAA
TAAAAACTTTGTTAATGGAAGAGAGTTCTTTTAGTGAAAAGGATGGTAAGTGGCTTCATGAG
CTTGCTCAGCACAACACATCTCTTGAGGTTTTAAACTTCTACATGACGGAGTTTGCCAAAAT
CAGTCCCAAAGACTTGGAAACCATAGCTAGAAATTGCCGCTCTCTGGTATCTGTGAAGGTCG
GTGACTTTGAGATTTTGGAACTAGTTGGGTTCTTTAAGGCTGCAGCTAATCTTGAAGAATTT
TGTGGTGGCTCCTTGAATGAGGATATTGGAATGCCTGAGAAGTACATGAATCTGGTTTTTCC
CCGAAAACTATGTCGGCTTGGTCTCTCTTACATGGGACCTAATGAAATGCCAATACTATTTC
CATTCGCGGCCCAAATCCGAAAGCTGGATTTGCTTTATGCATTGCTAGAAACTGAAGACCAT
TGTACGCTTATCCAAAAGTGTCCTAATTTGGAAGTTCTCGAGACAAGGAATGTAATCGGAGA
TAGGGGTCTAGAGGTCCTTGCACAGTACTGTAAGCAGTTGAAGCGGCTGAGGATTGAACGCG
GTGCAGATGAACAAGGAATGGAGGACGAAGAAGGCTTAGTCTCACAAAGAGGATTAATCGCT
TTGGCTCAGGGCTGCCAGGAGCTAGAATACATGGCGGTGTATGTCTCAGATATAACTAACGA
ATCTCTTGAAAGCATAGGCACATATCTGAAAAACCTCTGTGACTTCCGCCTTGTCTTACTCG
ACCGGGAAGAAAGGATTACAGATCTGCCACTGGACAACGGAGTCCGATCTCTTTTGATTGGA
TGCAAGAAACTCAGACGATTTGCATTCTATCTGAGACAAGGCGGCTTAACCGACTTGGGCTT
AAGCTACATCGGACAGTACAGTCCAAACGTGAGATGGATGCTGCTGGGTTACGTAGGTGAAT
CAGATGAAGGTTTAATGGAATTCTCAAGAGGCTGTCCAAATCTACAGAAGCTAGAGATGAGA
GGTTGTTGCTTCAGTGAGCGAGCAATCGCTGCAGCGGTTACAAAATTGCCTTCACTGAGATA
CTTGTGGGTACAAGGTTACAGAGCATCGATGACGGGACAAGATCTAATGCAGATGGCTAGAC
CGTACTGGAACATCGAGCTGATTCCATCAAGAAGAGTCCCGGAAGTGAATCAACAAGGAGAG
ATAAGAGAGATGGAGCATCCGGCTCATATATTGGCTTACTACTCTCTGGCTGGCCAGAGAAC
AGATTGTCCAACAACTGTTAGAGTCCTGAAGGAGCCAATATGA

**> COI1-like_MA_10427099g0010**

ATGCAGAGGCCGTGGAGGACGTCTGCGTGCGATGCGATTCCGGACGAGGTTTTGGAGTACGT
GATGGGGAACCTGGAAGACCCGCGGGATCGCAGCGCGGTTTCCCTGGTCTGCAAGAGCGCAC
GGTTCCCCAAGCTTGAGTCGCTGAAATTGAAGGGAAAGCCCAGGGCTGCCATGTTTAATTTG
ATTCCCCAGGATTGGGGCGGATACGCGAAGCCGTGGATTAACGAGATTTCCCAGAAATTTCT
CTGCTTAAAGGCTCTTCATTTGCGTAGAATGATCGTTACAAATGAGGATCTCGGGGTTCTGG
CTTGTGCCCGCGGCCACATTCTGCAGGTTCTTAAGCTGGAGAAGTGCTCGGGGTTTTCGACT
CTCGGGCTTCTCGAAGTCGCACGGTCCTGCAGGTAA

**> COI1-like_MA_10428988g0010**

ATGGAAATCCAGCGAGCGATGACGCTGGGCAACGAAATGGCAGAAGAAGCCCTGGAGTGCGT
GATGGATTACGTCAATGATCCCCGGGATCGAAGCGTGGTCTCTCAGGTGTGCAAGCAGTGGT
ACATGATCGATGCACTGACCCGAAAGCACGTGACCGTGGCCTTCTGCTACACAATCAAGCCC
GCCGATCTCACTCGGCGCTTCAGGCGGTTGGAGTCGTTGAAGCTGAAGGGGATGCCGAGGGC
CGCTATGTTCAACCTCATCCCAGAGGACTGGGGCGCCTACGCCTCCCCTTGGATTGAAGAGA
TCTCCTCTTCCTGCCTCTGCCTCAAGTCCCTCCACTTGCGGAGAATCATCGTCAAGGACGCC
GACCTCGCCACGCTCGTCGGAAGCCACGGTCAGATGCTGCAGGCCCTCAAGCTCGAGAAGTG
CTGCGGGTTCTCCACTCTCGGACTTCTGGAGATCTCTCGCTCTTGCAGATCTCTTAAGGTCT
TGTTTTTAGAGGAGAGTGAAATAAAAGATGAAGGTGGGGAATGGCTGCATGAGCTTGCTCTG
AGGAACATTTCATTGGAAGTGTTGAATTTTTATGTCCCATACTTAGAAAGCATTAACATGAG
AGATCTTGAGTTAATAGCCACGAACTGTCGAGCATTAACCTCTTTAAGGGTTAATGAATGTG
ATATTCTGGAATTAAGAGGTGTTCTGGATAAAGCTACAGAATTGGAAGAGTTTGGCGGTGGG
TCATTTCAATTTGTTAACAGTGATGAACATGCATTAGAAACTGATACATATCGAAGTGTTAA
ATTTCCTTCAAAGGTGATATCACTTATAGGACTCAATGACCTGAGCGAAACTGGGTTGCCTT
TTATACTTCCACGGGCTTGTAACCTGAAGAAATTGGATTTACAGTTTACATATTTGAGCACA
GAAAGTCATTGCGAGTTACTTTGTCTGTGCACTGGTCTTGAAGTTCTAGAGGTAAGGAATGG
AATTGGAGACAGAGGGTTAGAAGTTGTTGCAAACCACTGCAAAAAGTTAAGAAGACTGAGAG
TAGAGCCTGGGGAAGAAGAAGGTGAAGGTGGTGAGCAAAGTTCAGTCTCTCACAGAGGGCTT
TCTACCGTAGCTCAAGGCTGCCTCAATCTGGAGTTTATTGCTGTTTATGTTACAGATATAAG
CAACTTGGCATTGGAGACTGTTGGAGAATATTGCAAAAATCTTAGGGACTTTCGTTTAGTTT
TGTTGGATAAAGAAGAACAAATCACTGATTTACCACTTGATAATGGAGTCATGGCTTTGTTG
CGTGGGTGTAGCAAGTTGAACAGATTTGGCTTGTACTTAAGGCATGGGGGATTGACAGATAC
AGGACTTGGTTATATTGGGAAATACAGTGCAAATATAAGGTGGATGCTCTTGGGTTTTGTTG
GAGAATGTGACCAGGGTCTCCTTGAATTTTCAAAAGGATGCTTAAAACTGGAGAGACTTGAA
ATGAGAGGTTGTTGTTTCAGTGAATCTGCAATAGCGGCTGCAGTGCAGAACTTGAAATCATT
GAAGTATATCTGGGTGCAGGGATACAAAGGAACGACAACAGGTGAGAAACTTCTAGCTATGT
CACGACCATTTTGGAACATAGAGTTTACCCCACCTCTCAGAGAGACAGGGGATGAACTTGAT
GATATGGAAGAAGCAGTGATTGGACACCAACCAGCACAGATCTTGGCTTACTATTCTCTGGC
CGGAAGAAGAACAGATAATCCAAAATCTGTGATCCTTTTGTCGCCTCATTCTTGA

**> COI1-like_MA_10436355g0010**
ATGCTTGCATGTAAACAGGTTATGAATGTGACTGGAGATAGAGGTTTAGAAGTAGTTGTTGA
GAACTGTAAAAAATTAAGGCGACTTAGAGTGGAGCGTGGAGAAGATGAAGTTGGTTTGGAGG
ATGAGAAAGGCTCTGTTTCTCACAAAGGGCTCTCAGTTATGGCTCAAGGTTGTCCCAGTCTA
GAGTACATTGCTATGTATGCTTCAGATATGACTAACTCAACCTTAGAATCTGTTGGTAAATT
TTGCAAAAATCTGAGGGATTTTCTGCTAGTCTTGCTAGACAAGGAAGAACAAGTGACTGACC
TCCCACTGGACAATGGTGTCATGGCACTGCTTCTTGGGTGCCAAAAATTGAGGAGGTTTGGA
TTTTACCTAAGGCCTGGAGGATTGACGCACATAGGCCTTGGCTACATTGGAAAGTTTAGTAG
CAATGTGAGGTGGATGATTCTTGGTTATGTTGGAGAAACTAACATTGGACTTCTTGAGTTCT
CGAAGGGATGCCCAAATTTGGAGAAACTTGAATTAAGGGGTTGTTGCTTCAGTGAATATGCA
TTGTCCGTGACAGTTCTTAGCTTAAGGTCTCTAAAATATAGATGGGTTCAGGGTTACAATTC
AACGCCAACTGGAGTTGATCTTCTAGCTATGGAGTGTCCATTTTGGAACATAAAGTTTACTC
CAGCTTTCCATGTGACAGTGGATGGGTTTAATTTGGAAGCAGAAATTGTAGAGAATCCAGCA
CATATATTCTCTTATTATTCGCTAGCGGGAAGACGAACAGACCATCCAAATTCAGTAATTCC
GTTGACCTTATCATCATGGAATCGTCAGAATGTATATGGATATTAA

**> COI1-like_MA_108477g0010**
ATGGCCATGAAGCGGGCGGGATATGGATGCATATCAGAGGAGGCCCTGGAATGCGTCATGGG
TCAGCTGGAGGACCCGAGAGACCGGGGCTCGGTCTCTCTGGTCTGCAAGAAGTGGTACGACG
TGGATGCCTTCACGAGGAAGCATGTGACTGTGGCCTTCTGCTACTCGATACACGCCAGGGAC
CTCACCCGCAGGTTCACCAGGCTGGAGTCGCTCACGGTCAAGGGGAAGCCCAGGGCGGCCAT
GTATAATCTGCTTCCTGACGATTGGGGGGGTTATGCCAAGCCCTGGATAGACCAGATCTCGC

ACACCTGTCTCTGCCTCAAGACGCTCCATCTGCGCAGAATGATTGTTACCGATGATGATCTC
GCCACTCTCGTCAGGGGCCGCGGTCACATGCTGCAGGAGCTCAAGCTCGAGAAGTGCTCCGG
GTTCTCTACCAGGGGGCTCGAGGAAGTGGCGCACGGTTGCAGATCTCTTAAGACCTTGATGC
TGGACGAGAGTCAAATTGAAGAGGAAAGCGGGGATTGGCTACATGAGCTTGCTCTTAACAAT
TCTTCTTTGGAAGTGTTGAACTTCTACATGACAACAGTAGAAATGATCAATACCAGTGATCT
TGAGCTAATAGTAACAAACTGCCCCTCACTGACATCCTTAAAGGTTGGTGACTGTGATATAC
TGGATATGAGAGGTGTTCTGAGTAAGGGCACTGCATTGGAGGAGTTTGGTGGTGGTACATTT
AACACCAGTGAAGAGCATCCGACAGGGACCAATATGTCCCAGATGATTAAATTTCCTCCAAA
GTTGACATCATTGCTAGGACTAAACTTCATGATGGAGGCTGACATGCCTGCTATATTCCCAA
GAGCTTCTGCCCTTAAGAGATTGGATCTGCAGTACACATTTTTGAGCACAGAAAATCACTGT
CAGTTGGCAGGGCTCTGTCCTAATCTTGAAATTCTCGAGGTTAGAAATGTGATTGGAGACAA
AGGGTTAGAAGTTGTTGCAAATACTTGCAAAAAGCTGAAAAGACTTAGAGTGGAACGAGGAG
CAGATGACCCAACTTTGGAGGACGAACAAGGTTGGGTTTCTCACAAAGGGCTTTCCTCAGTA
GCTCAAGGCTGCCCCCTTCTTGAGTACATTGCTGTCTATGTTTCAGATATATGCAACTCAAC
ATTGGAGACTGTTGGTCAATGTTGCAAAAATCTCAAGGATTTCCGGTTGGTCTTGTTAGATA
AAGAAGAACACATCACTGATTTACCACTGGACAATGGAGTCATGGCTTTGCTACGTGGATGC
CAAAAACTGAGTAGGTTTGCATTTTATGTAAGGCCTGGAGGGTTGACGGATACAGGTCTTGC
TTATATTGGTGAGTACAGCACTAATGTCAGGTGGATGCTTCTAGGTTTTGCTGGTGAAACTG
ACCAAGGTATTCTCGAGTTTTCCAAGGGCTGCCCAAAGCTGGAAAGGCTAGAAATTAGAGGT
TGTTCTTTTAGTGAATCTGCATTGGCAGCTGCAGTGCTTCGTCTGAAATCGCTAAAGTACAT
ATGGGTTCAAGGATATAATGCAACTGTTACTGGCGCTAACCTTCTAGCTATGGCTCGACCTT
ATTGGAACATAGAGTTTTCTCCTGGTTTGCAATCGACAAAAGATGTGCTCGTTGAAGATATG
GCAGCAGAAAAAAATGCAGGATCGGGTAGCACAACTTTTGGCCTACTATTCTCTTGCTGGAAA
TAGGACAGATCACCCAGAGTCTGTAATTCCCTTAGCCTCACCTTTCCGGAATTACCAACAAG
TAGCTGTTTTCTAA

**> COI1-like_MA_15205g0010**
ATGTTAATAGCCACAAACTGTCCAGCATTAACCTCTTTAAGGGTTAATGAATGTGATATTCT
GGATTTAAGAGGTGTTCTGGAGAAAGCTACAGAATTGGAAGATTTTGGTGGTGGGGCATTTG
ATAACATTGATGGACATGCATTAGAAACTGAATATCAAAGTGTTAAATTTCCTCCAAAGGTG
ATATCACTTGTAGGACTCAATTACATGAGTGACACTGGGTTGCCTTTCATTCTTCCTCTGGC
TTGTAACCTGAAGAAATTGGATTTACAGTTCACGTTTTTGAGCATGGAAAATCATTGCGAGT
TACTTCGTCTATGCACTAATCTTGAAGTTCTAGAGGTACGGAATGTGATTGGAGACAGAGGG
CTAGAAGTTGTTGCAAACCACTGCAAAAAGTTAAGAAGACTGAGAGTAGAGGCCGGGGAAGA
AGAAGATGAAGATGAAGAGCAACATTTAGTCTCTCACAGAGGGCTTTCTACAATAGCTCAAG
GCTGCCTCAATCTGGAGTTTATTGCTGTTCATGTAACAGATATAAGCAACTTGGCATTGGAG
GCTGTTGGAGAATACTGCAAAAATCTTAGGGACTTTCGTTTAGTTTTGTTGGATAAAGAAGA
ACAAATTACTGATCTGCCACTTGATAATGGAGTCATGGCTTTGTTGCGTGGGTGTAGCAAGT
TAAACAGATTTGGATTGTACTTAAGGTCTGGAGGGTTGACAGATACTGGACTTGGTTATATT
GGGAAATACAGTACAAATGTAAGCAGCTGCAGTGCTGAACTTGAAATCATTGAAGTATATCT
GGGTGCAAGGATACACGAGATCGGTCTCGGCATTCCTGCTGAACCGCTCTTCCGATCTCACG
AATCTGCAATAGCAGCTGCAGTGCTGAACTTGAAATCATTGAAGTATATCTGGGTGCAAGGA
TACCAAGGAACAGAAACAGGTAAGAAACTTTTGGCTATGTCACGACCATTTTGGAACATAGA
GTATATCCAACCTCTCCCAGGGACAGGGGATGAACTTAATGTTATGGAAGAAGAATTGATTG
GACACCAACCAGCACAGATCTTGGCATACTATTCTCTGGCCGGAAGAAGAACAGATAATCCA
AAGTCTGTGATCCTTTTATCACCTCATTCTTGA

**> COI1-like_MA_263909g0010**
ATGGGCAACGAAATCGCAGAAGAAGCCCTGGAGTTCGTGATGAGCTACGTCAACGACCCCAG
GGACCGAAGCGTGGTCTCTCAGGTGTGCAAACAGTGGTACAGGATCGATGCCCTGACGCGGA
AGCACGTGACCGTGGCCTTCTGCTACACCATAAGGCCCGCCGATCTCACTCGGCGCTTCAAA
CGGCTGGAGTCGCTGAAGCTGAAAGGGAAGCCCAGAGCCGACATGTTCAGACTCATCACGGA

GGACTGGGGCGCCTACGCCCGCCCCTGGATTACCGAGATCTCCTCCTCCTGCCTCTGCCTCA
AGACCGAGATCTCCTCCTCCTGCCTCTGCCTCAAGTCTCTCCATCTCAGGAGAATGGTCGTC
AAAGACGACGACCTCACTATGCTCGTCCGGAGCCACGGTCACATGCTGCAGGCTCTCAAGCT
CGAGCGGTGCTCGGGCTTCTCCACTCTTGGACTTCTGGAAATCGCTCGCTCTTGCAGGTAA

**> COI1-like_MA_372199g0010**
GTACGAAATGTGATTGGAGACAGAGGGTTAGAAGTTGTTGCGGACCACTGCAAAAAGTTAAG
AAGACTGAGAGTAGAGCCTGGGGAAGAAGAAGGTGAAGATGAGCAAAGTTTAGTCTCTCACA
GAGGGCTTTCTACCATAGCTCAAGGCTGCCTCAATCTGGAGTTTATTGCTGTTTATGTAACA
GATATCAGCAACTTGGCATTAGAGACTGTCGGAGAATATTGCAAAAACCTTAGGGACTTTCG
TTTAGTTTTGTTGGATAAAGAAGAGCAAATTACTGATCTACCTCTTGATAATGGAGTCATGG
CTTTGTTGCGCGGGTGTACCAAGTTAAACAGATTTGGATTGTACTTAAGGCCTGGAGGATTG
ACAGACACAGGGCTTGGTTATATTGGGAAATACAGTACAAATGTAAGGTGGATGCTGTTGGG
ATTTGTTGGAGAATGTGACCAGGGTCTCCTTGAACTTTCAAAGGGATGCTCAAAACTGGAGA
GGCTTGAAATGAGAGGTTGTTGTTTCACTGAATCTGCAATAGCTGCTGCAGTGCTGAACTTG
AAATCATTGAAGTATATCTGGGTGCAAGGATACAAAGGAACGGCAACAGGTGAGAAGCTTCT
GGCTATGTCACGACTGTTTTGGAACATAGAGTTTACCCCACCCCTCATAGTGACAAGAGATG
AACTTGATGATATGGAAAAAGAATTGAGTAGACTCAAACCGGCACAGATCTTGGCTTACTAT
TCTTTGGCTGGAAGAAGAACAGATAATCCAGAGTCTGTGATTCTTTTATCACCTCATTCTTG
A

**> COI1-like_MA_437243g0010**
GTCAAGCAAGCACTTCACAGAAGAAGAAGAATGCAGAAGCCGTGGAGGACATCAGCAAGCTG
TGCGATTCCGGACGAGGTTCTGGAATGCGTGATGGGGTACCTGGAGGACCCCCGGGATCGCA
GCGCGGTTTCCCTGGTCTGCAAGCGGTGTCTTCCGATCTCTGCGCACGGTTCTCCGGGCTTG
AGTCGCTGAAATTGAAAGGGAAGCCCAGGGCTGCCATGTTTAATTTGATTCCTCCGGACTGG
GGCGGATATGCGGGGCCGTGGATTAACGGGATTTCTGAGACATTTCTCTGCTTAAAGGCTCT
TCATTTGCGCAGAATGATCATTACGGACGAGGATCTCAGGGTTCTGTCTCGCGGCCGCGGCC
ACATTCTGCAAGTGCTTAAGCTGGAGAAGTGTTCGGGGTTTTCGACTGTCGGACTCCTCGAT
GTCGCACGCTCCTGCAGGTAA

**> COI1-like_MA_63147g0010**
ATGCAGAGGTCGTGGAGGATGTCTGTGAGCGATGCAATTCCCGACGAGGTTCTGGAGTGCGT
GATAGGGAACCTGGAGGATCCGCGCGATCGCAGCGCCGTTTCCCTGGTGTGCAAGAGGTGGT
ACCGCGTAGATGCTCTCACTCGCAAGCATGTTACCATCGCGTTCTGTTACACCGTAAGCCCT
TCGTATCTCAGCGCTCGGTTCCCACGGCTTGAGTTGCTGAAACTGAAGGGGAAGCCCAGGGC
TGCCATGTTTAATTTGATTCCACCAGACTGGGGCGGATATGCGGAGCCGTGGATTAACGAGA
TTTCCCAGACATTTCACTGCTTAAAGTCTCTTCATTTGCGCAGAATGATCGTTACCGATGAG
GATCTTAGGGTTCTGGGTTGCGGCCGCGGCAACATTCTTCAGGTGCTTAAGCTGGAGAAGTG
CTCCGGATTTTCAACTCTCGGGCTCCTCGAAATCGCACGCTCCTGCAGGTAA

**> COI1-like_MA_63733g0010**
ATGGAGAACCAGCGGGTGCTGATGATGGACAGAGGAATGCCAGAAGAAGCCCTGGAGTGCGT
GATGAGCTACGTGAACGACCCCCGGGATCGAAGTGTGGTCTCTCAGGTGTGCAAGCAGTGGT
ACATGATCGATGCCCTGACCCGGAAGCACGTGACCGTGGCCTTCTGCTACACAATCAAGCCC
GCTGATCTCACTCGACGCTTCAAGCGGTTGGAGTCGCTGAAGCTGAAGGGGAAGCCGAGGGC
CGCTATGTTCAACCTCATCCAGGAGGACTGGGGCGCCTACGCCCGCCCTTCAGGAGGACTGG
GGCGCCTACGCCCGCCCTTGGATTGA

**> COI1-like_MA_646265g0010**
ATGGCACATGAATGCTTTTGGCATGCATGTGCCCTCTATCTCTTGCTCCTCCATAAATGCCC
TTGCGGCATTGTGCACTCATTTTACTGTATGGAACTTCTGAAGGAGGGCATACTGCTGCAGC

TGTCTACCCTACTAGAAGAGGAAGTTATGATAGCTGGATCACAGTCCGAGCTACACATAGGT
AAAAACTACAGACCTAACTCCAAAATCATATCTCTTAAGGTCTTGCTATTAGAGGCCAGTAC
GATAAAAGATGAAGGTGGGGAATGGCTACATGAGCTTGCTCTGAGAAACTCTTCATTGGAAG
TGTTGAATTTTTATGAAACATTGTTAGACAGCATCGACATGAGAGATATCGAGTTGATAGCC
ACGAACTGTCGAGCATTAACCTCTTTAAGGGTTAATGAATGCTATATTATGGATTTAATAGG
TGTTCTGGAGAAAGCTACAGAATTGGAAGAGTTTGGTGGTGGGTCATTTGTTAACAGTGAAG
CACATGCATTAGAAATTGATAGATATCAACTTGTTAAATTTCCTTCAAAGGTGATATCACTT
ATAGGACTCAATTTTATGAGTGGGTTGCCGTTTATACTTCCACGGGCCTGTAACCTGAAGAA
ATTGGATTTACAGTTTACTCTTTTAAGCACAGAAAATCATTGCGAGTTACTTCGTCTCTGCA
CTAATCTTGAAGTTCTAGAGGTATGTCTCTGA

**> COI1-like_MA_8264261g0010**
GTTACGAATGTGATTGGAGATAGAGGTTTAGAAGTAGTTGTTGAGAACTGTAAAAAAGTAAG
GCGACTTAGAGTGGAGCGTGGAGAAGATGAAGTTGGTTTGGAGGATGAGCAAGGCTCTGTTT
CTCACAAAGGGCTTTCAGTTATAGCTCAAGGCTGTCCCAGTCTAGAGTACATTGCTATGTAT
GCTTCAGATATGACTAACTCAGCCTTAGAATTTGTTGGTAAATTTTGCAAAAATCTGAGGGA
TTTTCGGCTAGTCTTGCTAGACAAGGAAGAACAAGTGACTGACCTCCCACTGGACAATGGTG
TCATGGCACTACTGCTTGGGTGCCAAAAATTGAGGAGGTTTGGATTTTACCTAAGACCTGGA
GGATTGATGGACACAGGCCTTGGCTACATTGGAAAGTTTAGTAGCAATGTGAGGTGGATGAT
TCTGGGTTATGTTGGAGAAACTGACATTGAACTTCTTGAGTTCTCGAAGGGATGCCCAAATT
TGGAGAAACTTGAATTAAGGGGTTGTTGCTTCAGTGAATATGCATTGTCCGTGGTAGTTCTT
AGCTTGAGGTCTCTAAAATATATCTGGGTTTAG

**> MYC2_AT1G32640**

ATGACTGATTACCGGCTACAACCAACGATGAATCTTTGGACCACCGACGACAACGCTTCTAT
GATGGAAGCTTTCATGAGCTCTTCCGATATCTCAACTTTATGGCCTCCGGCGTCGACGACAA
CCACGACGGCGACGACTGAAACAACTCCGACGCCGGCGATGGAGATTCCGGCACAGGCGGGA
TTTAATCAAGAGACTCTTCAGCAACGTTTACAAGCTTTGATTGAAGGAACACACGAAGGTTG
GACCTACGCTATATTCTGGCAACCGTCGTATGATTTCTCCGGCGCCTCCGTGCTCGGATGGG
GAGATGGTTATTACAAAGGTGAAGAAGATAAAGCAAACCCGAGACGGAGATCGAGTTCGCCG
CCGTTTTCTACTCCGGCGGATCAGGAGTACAGGAAAAAAGTGTTGAGAGAGCTTAACTCGTT
GATCTCCGGTGGTGTTGCTCCGTCGGATGACGCTGTTGATGAGGAGGTGACGGATACGGAAT
GGTTTTTCTTGGTTTCGATGACGCAGAGCTTCGCTTGCGGTGCGGGATTAGCTGGTAAAGCG
TTTGCAACGGGTAACGCGGTTTGGGTTTCCGGGTCAGATCAATTATCCGGGTCGGGTTGTGA
ACGGGCTAAGCAAGGAGGAGTGTTTGGGATGCATACTATTGCGTGTATTCCTTCGGCGAACG
GAGTTGTGGAAGTCGGGTCAACGGAGCCGATCCGACAGAGTTCGGACCTTATTAACAAGGTT
CGAATTCTTTTCAATTTCGACGGCGGAGCTGGAGATTTATCGGGTCTTAATTGGAATCTTGA
CCCGGATCAAGGTGAGAACGACCCGTCTATGTGGATTAATGACCCGATTGGAACACCTGGAT
CTAACGAACCGGGTAACGGAGCTCCAAGTTCTAGCTCCCAGCTTTTTTCAAAGTCTATTCAG
TTTGAGAACGGTAGCTCAAGCACAATAACCGAAAACCCGAATCTGGATCCGACTCCGAGTCC
GGTTCATTCTCAGACCCAGAATCCGAAATTCAATAACACTTTCTCCCGAGAACTTAATTTTT
CGACGTCAAGTTCTACTTTAGTGAAACCAAGATCCGGCGAGATATTAAACTTCGGCGATGAA
GGTAAACGAAGCTCCGGAAACCCGGATCCAAGTTCTTATTCGGGTCAAACACAATTCGAAAA
CAAAAGAAAGAGGTCGATGGTTTTGAACGAAGATAAAGTTCTATCATTCGGAGATAAAACCG
CCGGAGAATCAGATCACTCCGATCTAGAAGCTTCCGTCGTGAAAGAAGTAGCAGTAGAGAAA
CGTCCAAAGAAACGAGGAAGAAAGCCAGCAAACGGTAGAGAAGAGCCACTAAACCACGTCGA
AGCAGAGAGACAAAGACGCGAGAAACTAAACCAAAGATTCTACGCGTTACGAGCGGTTGTAC
CAAACGTTTCAAAAATGGATAAAGCTTCGTTACTCGGTGACGCAATCGCTTACATCAACGAG
CTTAAATCCAAAGTAGTCAAAACAGAGTCAGAGAAACTCCAAATCAAGAACCAGCTCGAGGA
AGTGAAACTCGAGCTCGCCGGAAGAAAAGCGAGTGCTAGTGGAGGAGATATGTCGTCTTCGT
GTTCTTCGATTAAACCGGTGGGGATGGAGATTGAAGTGAAGATAATTGGTTGGGACGCAATG
ATTAGAGTTGAATCTAGTAAGAGGAATCATCCGGCGGCGAGGTTGATGTCGGCGTTGATGGA
TTTGGAGTTGGAAGTGAATCACGCGAGTATGTCGGTGGTTAACGATTTGATGATTCAACAAG
CGACGGTGAAGATGGGTTTTAGGATCTATACGCAAGAACAGCTCAGAGCAAGTTTGATTTCA
AAAATCGGTTAA

**> MYC2-like_MA_15962g0010**

ATGATGCAGTCTTTTGTTAATGAATGCGGGGTGCCTGGGAAGGCATTTTCCAGTGCCATGCC
TGTTTGGATTATCGGCTCAGAAAGGCTTCAGGGCTACAACTGTGACCGCGCTCGTCAGGCTC
AGCAATTTGGCATTCAGACCATGGTATGTATTCCGACACTTAACGGAGTTGTTGAGTTGGGT
TCCACGGATTTGATCCCCCAGAACTGGGATTTGATACAGAAGGCTAGAGATTCGTTTACTTT
TACGCTTCCGGACACTGCTTTGTGGGAAGAAAATCATACCCAAAATGATCCTGACCCGGCTC
TTTGGTTGACTGAACCCCCGGCTGAGCCTAAAACAGAAACTGAAAAGAAACAACAGCAGATA
CCCAATGCGGAAACTGAAGCTCTTCACAGCTTTTTTCCTCATGAATTAGGGTTTTCTGACCT
GGGGTTTCTGAATGGCGAGGGAGAAAACGCGTCACAGAAGCTTGTGGTTGAGGAGTGTGGGC
AGAAAGAAGATAAATCATCTCAGGGCTTTACTGGTTCTCAGGTTTCGTACCAACAGAACTGG
CAGGCTCAGACGACCTGCAAGACTGAAGTTGTGGATATTCCGGTGTTTCAACCTGGGAAAAG
GACTAATACTAATGGGATTACTCTGAGCTTTGAAAACCCCTACGGTGCTCAGGGTTTAGTGA
GTTGGGGGGATGAGAAGATGAAGAGGTCTGTGAGGAATGGGAATGAAGATGGTGGGAGTGGT
TTGTGTTTTTCTTCTGAAGTTACTGCTGTTTCTGCTTCTACTTCTGCTACTGCTGCTGCTGC
TGCTTTGCCTGTGAACGGTGCTGCTGGGGTTTCGTACCAACAGAACTGGCAGGCTCAGACGA
CCTGCAAGACTGAAGTTGTGGATATTCCGGTGTTTCAACCTGGGAAAAGGACTAATACTAAT
GGGATTACTCTGAGCTTTGAAAACCCCTACGGTGCTCAGGGTTTAGTGAGTTGGGGGGATGA
GAAGATGAAGAGGTCTGTGAGGAATGGGAATGAAGATGGTGGGAGTGGTTTGTGTTTTTCTT
CTGAAGTTACTGCTGTTTCTGCTTCTACTTCTGCTACTGCTGCTGCTGCTGCTTTGCCTGTG

AACGGTGCTGCTGGGGTGAGATCTAGTGTTGAATCGGAGCATTCAGATATAGAGGCGTCTTT
TAAAGAGGCCGAATGCAGCCAGGCCATTGTTGAAAGGAGGCCTCGGAAACGGGGAAGGAAGC
CTGCCAATGGTAGAGAAGAACCTCTGAATCATGTAGAAGCTGAAAGGCAGAGGCGAGAAAAG
TTGAACCAGAGGTTTTATGCACTCCGTGCTGTGGTTCCCAATGTGTCCAAGATGGATAAGGC
TTCTCTGTTGGGTGATGCCATTTCTTACATTAATGAGCTCCGAAACAAGGTGCAGGATTCCG
ATTCTCATAAAAAGGACTTGCAGGCTCAACTCGAGGCCTTGAAGAAGGAATTGGTAGCCAGA
GAATCCGTTGCTTCTGGATTTTCTGGCAGTAATTTTGGTTTGCTAAAAAACCCATCTGCTGC
TGATCCTTCAAACCTTGATGTCAAAGGCTTTGGTTTGAAAAACCAGTGTCCTAATATTGAGC
TTGAAGTACGAATCCTTGGTCGAGAGGCCATGGTCAGAGTTCAGTGTCCCAAGCAGAACCAT
CCTGTTGCAAGATTGATGGTTGCATTTAAAGAGCTTGAACTTGAAGTCCATCATGCCAGTGT
ATCCACAGTGAAGGAGTTGATGATTCAAACGGTTATTCTGAATATGACAGGTATTGTATATA
CGCAGGAACAACTAAATGCTGCATTATTAAGGAAAGTAGCAGATCCTGGCCTTAGATAG

**> MYC2-like_MA_5838g0020**
ATGGTGGAGGCATTCATGGCGTCGTCTTCGTCGTCGTCGTTTTTGGCGGAGTATCCGCCATG
GAGCAACGCAGGCGGGGACCCGAACGGTTCATTCAATCAGGAGACACTGCAGCAGAGGCTGC
AGCTCTTGATCGACGGAGCGCGCGAGAGGTGGACATACGGGATTTTCTGGCAGTGGTCGTAT
GAGGCCGGCGGTGCCGTGGTCTTCTTCTGGGGCGACGGCTACTTCAAGGGCGCCCTCGAGGA
CGAGAGGACGAACACCGTGGCTGTCGCCAAGCGGACAACGAAGGAGAACGCGGCGCACCAGG
AGCTCAGGAAGAAGGTGCTCCGCGAGCTGCACGAGCTCATCGACGGGGCCTCCGATCCCGCC
GACGAGGAGGTGACGGACGTGGAGTGGTTTTATCTGGTCTCCATGACGCATTCGTTCACAGG
GGTGGAGGGGGTTCCGGGGCACGCGTTCATGTCCAGCGCGCCCGTGTGGCTGTCTGGGTCGC
TGAAATTGGAGTCGTTCGGATGCCAGAGGGCCAGGCAGGCTTCTCAGTTCGGGATTCAGACC
ATGGTATGTATTCCAACTCCTAATGGCGTTGTGGAATTGGGTTCTATGGATCTGGTCTGTGA
GAACTGGGGTCTGCTGCAGCAGGCCAAGAGCTCATTTACTTTTTCCTCTAGTTTCTGGGAAG
ATAACGGGAATGGTAATATTAATCATAACAATAACCATAATAATAATTATGGTACTAATAAC
CAGTCGTTATGGAATCCGGGGTCTCCATTCTTGACCCAGGAATCGATCTTGGGTGACTTGAG
TTTTCTCAATAACGAGGAAAGCCAAAACCGGAATTCGTCTGCTCAAAAATCCTTGTCCATCT
TGGAAGAAAACCGAAACCCTTTACCCCCTTTTACTGTTCAAAAGCAAGTAGCCCTGGAGGAA
AACCGGATTCCGCCCTCTTTTCCTGCTCAGAAATCTGCCGTCCTTGACGAGAGAACCTCCTT
ACCTCTTGTTCAGAAAACTGGCATCTCCGAAGGAGCTCATAATCCCCTGCCGTTCCTTTCAA
AGCCACCTCCTGGAGGCGGCTTTGATGAGAAGTCTAATACTTTGCCATATTCTGCTGTTCAA
AGGCCTGCTATCATTGATGAGAACTGCAATTCTTTGCTTTCTCAGAAATCCAGCATCTGCAG
TGAGATTCTTAATTCTTACCCATATGTGACCGGTCCAAAATCTGTTGATTTCAGCGAAACTT
GTAATTCTTATCCAGTTCCAGCTCTTCAGAAAACTGGCATTGAAAACAGTAATCAGAATTTA
CTACAATCCGCTGCTGTTCTGAAACCTAACATAGCGGACCAAATTGGTAATCCGAATTCAAA
CCCAAATCCGCTGCTTTTTTCTGTTCAGAAAACTTGTACTATTGATCAAAGCAAGGGATTCA
TCAAAGATCTCCTGATAGAGGACGACAAGCCCAAGCCCTTGTTGGATTTCCAGAGACACACT
CTGAGCTTTGCAAATGGATACTCGCAGAATATGGTGGAGGAGAAGATGGTGAAACCTCTCAG
TATTGATGATGAAAAGCCCAAGTCCTTGCCTACAATTTCCTCTGGTGCTGTCTTTGGTGGAG
TTCGATCTAGTATTGAATCTGACCATTCTGATGTAGAAGCTGCGTCCTTTAAGGAAGCAAGC
CAAGCTGTCATTGAGAAAAAGCCCCGGAAAAGAGGAAGAAAGCCTGCAAATGGTAGAGAAGA
GCCTCTGAATCATGTTGAAGCTGAGCGTCAACGGCGCGAGAAACTGAATCAGAGATTCTATG
CACTTCGCGCTGTAGTACCAAATGTGTCAAAGATGGATAAAGCTTCATTGCTTGGGGATGCT
GTGTCCTATATTAATGAACTTCAAAGCAGAGTTCAAGACATTGAATCTGAGAAAAAGGAGCT
TCAAGCCCAAATAGAAGCTACCAAGAAGGAATCCTCGTCTTCTCACTCTGCTTTCTCTGGTA
CCAATTTGGGATTCATCAAGGACCAGTCCGGTTCATCTCAAAAGCCTGATGTCAAACGATTT
GGCACAAAGGAATGCTCTGCACTAGATCTGGAAGTTCGAATCCTTGGTCCAGATGCCATGAT
CAGAATTCAGTCAGCTAAGAAGAATCATCCTGCAGCCAGATTGATGACATCACTGCAAGACC
TAGAGCTTGAAGTTCATCATGCCAGTGTCTCAACAGTCGACGAGTTAATGCTTCAAAATGTG
ATTGTCAAGTTACCAAGTAGTTTATATACTGAAGAACAACTCAATGCCATCCTGTTGAAGAA
ATTATCAGATCCAAAATTTAAGTAA

**> MYC2-like_MA_10435905g0020**
ATGGATACTTTAATGGCGTCATCAGTGGTAGATCAGAGGTTCAGCCAGGAAACATTGCAGCA
GCGCCTGCAGACTCTAGTAGAGACGGCCTCAATAGTGTGGACTTATGCGATTTTCTGGCAAG
TTTCTTATGAATCTAGCGGTGCCATTCAGCTATGCTGGGGAGACGGCTATTACAAGGGGTCG
AGGAATACGGAAGAGGATGAGCGGCTTAGGATGCGCAGCCGTTTGACCGTGAGCCCCGCCGA
CCAGGAGCTGAGGAAAAAGGTATTACGCGATTTGCATTCTATGATTAGCGGGAGCGATGAAG
GAAACCAGCAGGATAATAGTAGTGTTAGCGTAGATGAAGAAGTTACTGATGCTGAATGGTTT
TATTTAATTTCGATGATGCAATCCTTTTTATCTGGGTTTGGGGTACCCGGTACGGCATTTTC
TACCGGAGCTCCTGTTTGGATCGTCGGAGCGGAGAGACTGCGGGTATCGACATGCGACAGGG
CAAGGCAAGCTCATGACCTTGGGATTCAAACACTTGTCTGTGTTCCGATTCAGGGCGGCGTG
GTTGAATTTGGATCGACGGAAGACATTGTTGAAAACTGGTTGTTTCTGGAGCAGGTCAACCG
TTCGTTTAAATATAACCTCAACCAAACGCATGATAACCTGTTTCAAGTCCAGTCTTTATGGC
CAGAGGAAACTTTGAGTGTTAAAAGCAGCAATACTATGCAATCCGCGCCATGCATCGAGCCT
GTGAACAATGCTGAAATCCAGTCCCTGAACTCTGCTTTAGCTCGGGAATTACCAGTGACAGG
CAAACAAAAGGCATCTGTGTTCGCTGAACAGAGTTCTTTAGTGGTTAAAGACGACAAATCTT
TGCTGCATCCCCTAACTCAGCAGACCGAAGCCCTTGAAGCGCCGGCCATCCGTATTCCAGAA
ACAGTAAACGGTACAGAGCCACAAACCCGCGCATTGGGCTTTAAAGGCAGTGAAAAGAATGT
GATAAAGCCATCAATAAAAGAGGATACCATTGGTTTACTGTCTAATCCGCCTGGAATTGTCA
TAGGAGGCCTGCGTTCTAGCATTGAATCAGAGCTGTCTGATGCAGAACCCTCTGCCTCAATT
AAAGATTCAACTTCTGCTGTAGTCGAGAGAAAACCACGGAAACGTGGGAGAAAGCCTGCAAA
TGGCCGCGAAGAGCCTCTGAATCATGTTGAGGCTGAGCGGCAAAGGCGTGAGAAATTGAACC
AGAAATTTTATGAGCTTCGTGCTGTGGTTCCCAATGTGTCGAAGATGGACAAAGCTTCTCTG
CTCGGCGATGCCGCTGCTTATATCAAAGACCTCTGTTCCAAACAGCAGGATTTGGAATCCGA
GAGGGTTGAATTGCAGGATCAAATTGAGTCTGTAAAGAAGGAATTATTGATGAATTCTTTGA
AGTTGGCAGCTAAAGAAGCAACAGATCTTTCAAGTATTGACCTTAAAGGTTTTAGCCAGGGG
AAGTTCCCCGGCTTGAATTCAGAAGTTCGCATTCTTGGCCGAGAGGCGATAATAAGAATTCA
ATGCACTAAACATAATCATCCTGTTGCGAGACTGATGACAGCACTGCAAGAACTTGATTTGG
AAGTCCTCCATGCGAGTATTTCTACTGTGAAGGATTCGTTAATTATCCAGACAGTCATTGTT
AAAATGACCAGAGGTTTGTACACGGAAGAACAACTTCATGCGCTGCTTTAGCAGGTAATGGA
CTGCTCAAGGATTGTTTTAGAATTTGCATTGGTCTCAGCATAA

**> MYC2-like_MA_9725g0010**
ATGACAACGGAGGAGAGAGCAACACAACATGAGCTCAGGAAGAAGGTGCTCCACGAGCTCAT
CAATGGGGCCTCTAATCCTGCGGTCGATGAGGAGGTGACGGATGTGGAGTGGTTTTATCTAG
TCTCCATGATGCATTCATTCATAGGGGCCAAGGTGGTTTCGAGCCATGCAGTCATGTACAAT
GCACCTATATGGCTCTCCGGGTCACCGAAATTGGAGTCATTTAGATGTTAG

**> MYC2-like_MA_140086g0010**
ATGGGCAACGAAGGCGCGGGATCGATGCGGCTGTGGGGCGATGATAATAATGTCATGATCGA
GGCCTTCATGGGGAACCTGGATTACTCCTACTCGTTCCCCTGGAATGGCATCGACGCCAATC
CCTCCGCTCTACCCTCGCCCGCCCACTTCGCATCGTCTGCCGCCAGCGTTGCTATTGCTACG
CCCTTCAATCAGGACACGCTGCAGCAACGACTGCTGGCGCTTGTGGAGGGCGCTACCGAGAG
CTGGACG

**> JAZ10_AT5G13220**
ATGTCGAAAGCTACCATAGAACTCGATTTCCTCGGACTTGAGAAGAAACAAACCAACAACGC
TCCTAAGCCTAAGTTCCAGAAATTTCTCGATCGCCGTCGTAGTTTCCGAGATATTCAAGGTG
CGATTTCGAAAATCGATCCGGAGATTATCAAATCGCTGTTAGCTTCCACTGGAAACAATTCC
GATTCATCGGCTAAATCTCGTTCGGTTCCGTCTACTCCGAGGGAAGATCAGCCTCAGATCCC
GATTTCTCCGGTCCACGCGTCTCTCGCCAGGTCTAGTACCGAACTCGTTTCGGGAACTGTTC
CTATGACGATTTTCTACAATGGAAGTGTTTCAGTTTTCCAAGTGTCTCGTAACAAAGCTGGT
GAAATTATGAAGGTCGCTAATGAAGCAGCATCTAAGAAAGACGAGTCGTCGATGGAGACAGA
TCTTTCGGTAATTCTTCCGACCACTCTAAGACCAAAGCTCTTTGGCCAGAATCTAGAAGGAG
ATCTTCCCATCGCAAGGAGAAAGTCACTGCAACGTTTTCTCGAGAAGCGCAAGGAGAGATTA
GTATCAACATCTCCTTACTATCCGACATCGGCCTAA

**> JAZ10-like_MA_10229741g0010**
ATGTACCGCGCAGCTATGGACTCCATGGGCGTCGATACTAAGGGATCCAGTATTAACCGAGC
CCCTGAATCTGGTCCGGCGGATCAACTCAAAAACGATAGAGAAAGAGCCTGCATGAACAATC
TATCGTGGATAAAACCCCAAATGATGCGACAAGTGCTTTCGTTTAGAGATGAAGCAAGAAAC
GCGTCATGCCATGGAGTATCTCAGGTTCGTATGGATTCCTGCGCTCCAGGGCTAATGGGATT
ATTTCGTCAGCCAATAGGTTTTGTGGCTTCCATTTCTCCTGCTGCAAACCACGAGACCTCGG
CTTGCAAATATACTCATAATACCGCGGCAGCAGGTAGTGTGCAGTCACAACGATGGCCTTTC
CTCTCCCATGGATCTGTCCAGACAAATGGGACAATTTCCAGGCCGGCGTCTGCAACGGCTAC
AGAAGGCAAGACAACGACTGCCCCGTTGACGATCTTCTATAACGGCACGGTCAGCGTTTACG
ATGTCCCTGCGCACAAGGCTAAAGCAATCATGATGCTGGCCACAACTGCAATCTCTAATTGT
AAGGCAACTTCTCCTACCTCTACCATTACGACTACATCCGGAGCTTCAGCCAGTAAACTGAT
CTCCATGGCAGTACCAACCGAGACATTTACAGGAGCGGCCGCGCAGCAACAAACAGCTTGCA
AGCCGAATGCTGGACTGCCAATTGCGAGGAAACAATCGCTGCAACGTTTTCTCGAGAAGCGG
AAAGAGAGACTGAACATCGCCAGTCCGTACGCGATGAAAACCTCCGAATCCAACGAGGAATG
CGCCAGTGCACCCATCGTTTTTCCGACGGTGTAA

**> JAZ10-like_MA_8757791g0010**
ATGATGCGACAAGTGCTTTCGTTTAGAGATGAAGCAAGAAACGCGTCATGCCATGGAGTATC
TCAGGTTCGTATGGATTCCTGCGCTCCAGGGCTAATGGGATTATTTCGTCAGCCAATAGGTT
TTGCGGCTTTCATTTCTCCTGCTGCAAACCACGAGACCCCGGCTTGCAAATATACTCAGAAT
ACCGCGGCAGCAGGTAGTGTGCAGTCACAACGATGGCCTTTCCTCTCCCATGGATCTGTCCA
GACAAATGGGACAATTTCCAGGCCAGCGTCTGCAACGGCTACAGCAGGCAAGACAACGACTG
CCCCGTTGACGATCTTCTATAACGGCACGGTCAGCGTTTACGATGTCCCTGCGCACAAGGCT
AAAGCAATCATGATGCTGGCCACAACTGCAATCTCTAATTGTAAGGCAACTTCTCCTACCTC
TACCATTACGACTACATCCGGAGCTTCAGCCAGTAAACTGATCTCCATGGCAGTACCAACCG
AGACATTTACAGGAGCGGCCGCGCAGCAACAAACAGCTAACAAGCCGAATGCTGGACTGCCA
ATTGCGAGGAAACAATCGCTGCAACGTTTTCTCGAGAAGCGGAAAGAGAGACTGAACATCGC
CAGTCCTTACGCGATGAAAACCTCCGAATCCAACGAGGAATGCGCCAGTGCACCCATCGTTT
TTCCGACGGTGTAA

**> JAZ10-like_MA_15372g0010**
ATGCAAAGCATCTGGATATGTACAATCCCGTGCGAAGGTGGGAACTGGAACCAAGTGCCTGA
CCCTGCCTATGCTCGTCTTCGCCTCCTTTACCTTCAGCGCAGAGGGAGCCTTGATCTTGCTT
GGGCCTCGACCTCAAGGGCTCCAGTATTAACCGAAACCCTAAATCTGAACCTGCGGATCAGT
TCAACAATGACAAGGACAGAGAATCAGTCGTGGATAGAGCCCCAGGTGATTCAGCAAGTCCT
CTCGTTTACGGATAGGGCGAAAGGCGAGTCGTGCATCTATAGAGGGTCTCAGGTGCTGTTGA
ATTCCTCCGCTCCGGGCTTAATGGAGTTATTTCCTAAAGAAACAGTATTTGCAGCTTCCAGT
TCTGCTGCCACAAAGCACAAGAGCCCTTATTGCAAGAGTACTCTTCAAGAGACCGCAGTTCC
AGCTAGTGTGCAGTCACACCGATGGCCATTCCTCTCCCACCGATCTGTCGAGACAACTGCTA
CGATTTCGAGGACGGCCTCTGCCCCGTTGACAATCTTTTATAACGGGAAGGTCAGCGTTCAT

GATGTCCCTGCTCACAAGGCTGAAGCGATCATGATGTTGGCCACCACTTCAATTTCTAATCG
CCAGGCAAGTTCTTCTCCTCCCACTACCTATGGGCCTTCAAGCGGTACATTCAGGTCCACGG
AAATACCCGCAGAGCCATTCGAGGCCGCATCTGCGCAGCCAGAATCGACTCGAAAATTGATA
ATCGGAATGCCCGTTGCGAGGAAACAATCGCTGCAGCGTTTTATTGAGAAGCGAAAAGAGAG
ACTGAACAGCGCAAGCCCTAATTACAAAATGAATCCGACGACATCAAAACCCGTCGCGTGCG
TCGAGGGATGCGGCGGTGCTTCCCTTATTTCTCCGATGGTGTAA

**>AOC1_AT3G25760**
ATGGCTTCTTCTACAATCTCTCTCCAATCTATCTCCATGACAACTCTCAACAATCTCTCATA
TAGTAAACAATTTCATCGAAGCTCTCTTCTTGGTTTCTCTAAATCCTTCCAAAATTTTGGTA
TCTCATCTAACGGTCCAGGTTCCTCCTCTCCGACAAGTTTCACGCCCAAGAAGAAACTCACT
CCTACTCGAGCTCTCTCTCAGAACTTGGGAAATACCGAAAACCCCAGACCAAGCAAAGTTCA
AGAACTGAGCGTGTACGAAATCAATGATTTAGATCGACACAGCCCAAAAATTCTTAAAAACG
CATTCAGCTTTAGGTTTGGTCTCGGAGATCTCGTCCCATTCACAAACAAACTCTACACCGGC
GATCTCAAGAAACGCGTGGGAATCACGGCTGGTCTCTGCGTTGTGATCGAACACGTCCCTGA
GAAGAACGGTGATAGATTCGAAGCTACGTACAGCTTCTACTTCGGAGACTATGGCCACTTGT
CCGTACAAGGACCATACTTGACTTACGAGGATTCGTTCCTTGCCATCACTGGTGGCGCTGGA
ATCTTTGAAGGTGCCTACGGACAGGTTAAGCTTCAACAGCTTGTGTATCCGACAAAACTGTT
CTACACTTTTTATCTTAAAGGATTGGCTAATGATTTGCCACTGGAGCTCATAGGAACTCCGG
TACCACCGTCTAAGGACGTGGAGCCAGCACCGGAAGCTAAGGCGCTTAAGCCCAGTGGAGTT
GTAAGCAACTTTACAAATTAG

**> ACO1-like_MA_137604g0010**
TGGCAGCGTCTGCTCAAAGCCGAATACAATTTCACAGCTCCTCTCTGTTGAAGCCAGAGGCA
TCAATTCGACATCCTCAGTACAGATACTGGAAATGCTATTTCACAGCTCCATACAATTTCAC
AGCTCCTCTCTGTTGAAGCCAGAGGCATCAATTCGACATCGGGCGAAGGCCATCTCTTCTTC
TTCTTCTGCGCAAGGGCCTCCAAGCGGCTCTGTTCAGGAGTTGCACGTGTATGAGATAAATG
AATTGGACAGAGGGAGTCCTGCTTACCTTCGTCTCAGTCAGAAGCCCGTCAATCATCTGGGA
GATCTAGTACCATTTAGCAATAAGCTGTACACTGGCAATCTACAGAAGCGTATAGGAATAAC
AGCTGGGATCTGTATTCTCATCCAACATTTTCCTGAGAAGAAAGGAGACCGTTATGAAGCAA
TCTACAGTTTTTACTTTGGAGATTATGGCCACATTGCTGTGCAGGGACCTTATCTGACATTC
GAGGACGAGGAATCATACCTTGCAATCACTGGGGGTTCTGGCATATTCAACGGAGTTCGTGG
GCAAGTGATGTTAAAGCAGCTTGTCTTTCCTTTCAAGATATTCTACACATTTTACTTAGAGG
GCATTCCAAGCCTGCCGGAAGATCTGCTGGGTGAGCCTGTGGCACCGACACCTGATGTGGAG
CCATCTCCCGCTGCAAAGAATAAAGAGCCTCATGCCTGTGTGCCCAATTATACACAGTAA

**> ACO1-like_MA_56386g0010**
ATGGCTGCTACTGCCACTGCCACCTCGATCTCATTAAAATCATCCTTGGGAAACAACAAAAG
CCTGCATCAATTGGTTTGTTCAAACTCAGAGGGCAACAATACTAATTCTATAGGCTTCAAAC
CTGCTACTTGTTTTAAGCTGTTAAAGGGTGAGATTTTTGGGGTACAGTTGGCTACTGAGAGG
CAAATCCAGTTTGACAGCTCCTCTCTGGTGCGAGTGAGGGCAGGGGCAGAGACATCGATATC
AATTCGACGTCGAGTTAAGGCCAACGCTTCTTCTGCTCAAGGCCCCCCAAGCGGCTCTGTGC
AGGAGTTGTATGTGTATGAGATGAATGAGTTGGACAGAGGAAGCCCTGCTTACCTTCGTCTG
AGTCAGAAGCCCGTCAATCATCTGGGAGATATAGTGCCATTTAGCAATAAGCTGTACAGTGG
CAATCTGCAGAAACGTCTAGGTATAACAGCTGGGATCTGTATTCTCATCCAGCATTTTCCTG
AGAAGAACGGAGACCGTTATGAAGCAATCTACAGCTTTTACTTTGGAGATTATGGCCACATT
TCTGTGCAGGGGCCTTATCTGACATACGAGGAATCATACCTTACAATTACTGGGGGTTCTGG
CATATTCAGTGGAGTTCATGGGCAAGTGAAGTTAAAGCAGATTGTCTTTCCTTTCAAGTTAT
TCTACACTTTTTACTTGGAGGGCATTCCAAGCCTGCCCGAAGATCTGCTGGGTGAGCCTGTA
TCCCCAACACCGGATGTGGAGCCATCTCCCTCTGCAAAGAATGAAGAACCTCATGCCTGCCT
GCCCAACTATACTATGTAG

### Supplementary Table S1 Primers used for qPCR experiments in this study

| Gene ID | Given Name | Forward (5'-3') | Reverse (5'-3') | Efficiency |
|---|---|---|---|---|
| MA_108477g0010 | *PaCOI1* | GCAACTGTTACTGGCGCTAACCTTC | GCCATATCTTCAACGAGCACATC | 2,57 |
| MA_5838g0020 | *PaMYC2* | AATACTTTGCCATATTCTGCTGTTC | GGATAAGAATTACAAGTTTCGCTGA | 2,353 |
| MA_6326g0010 | *PaJAZ3* | AAGGGAGAGCAAGCAGCAGGAACAAC | AGAGGCTCCGACAACAGGCAAGAAAG | 2,32 |
| MA_10229741g0010 | *PaJAZ10* | TGGCTTCCATTTCTCCTGCTGCAA C | CCGGCCTGGAAATTGTCCCATTTGTC | 2,942 |
| MA_56386g0010 | P*aAOC1* | CTCCTCTCTGGTGCGAGTGA | GGGCTTCCTCTGTCCAACTC | 2,466 |
| MA_50378g0010 | *PaEIF4A1* | TTGGTCGGAGTGGACGATTTGG | GTCCTGCAGCATTCTCTCGTCA | 2,041 |
| MA_13020g0010 | *PaUBC28* | GGATCTCTGTTAACCGCGTCGTTG | AGGATCCGCTTAGACGCCATT | 1,989 |
